# POLQ variants with aberrant DNA polymerase activity protect against UV-induced cell death

**DOI:** 10.1101/2025.11.26.690880

**Authors:** Steven Weicksel, Corey Thomas, Ethan Hall, Kylie Davis, Chase Michalczik, Sreerupa Ray, Jamie Towle-Weicksel

## Abstract

DNA polymerases are important for maintaining genomic stability by protecting against mutagenic lesions caused by external and internal factors. If left unrepaired, DNA damage can lead to replication errors resulting in changes in DNA sequences that could impact peptide sequences and gene expression, as well as lead to chromosomal breaks. Highlighting the importance of DNA polymerase repair function, variant DNA polymerases and cofactors of DNA repair pathways have been identified in many different cancer types. Recently, variant forms of DNA polymerase Q (POLQ) have been identified in patient isolated melanoma tumors. Previous work identifying biochemical characteristics of these variants has shown that they display aberrant DNA polymerase activity compared to wild-type (WT). To better understand the role these variants have in DNA repair, genomic stability and cell survival, we tested their ability to bypass and extend DNA past cyclobutane pyrimidine dimers (CPD) as well as prevent cell death when exposed to ultra-violet (UV) radiation. Biochemically we show that the patient derived variants of POLQ tested here display decreased efficiency during DNA bypass and extension of CPD lesions and prefer to incorporate purines. In addition, two of the three variants protect against UV induced cell death. Together these data suggest that POLQ variants can support cell survival and further supports that POLQ variants can act as both protectors of cell viability as well as drivers of genomic instability, characteristics important in cancer cells.

**HIGHLIGHTS:**

▯ POLQ variants have decreased efficiency during bypass and extension of CPD damaged DNA compared to WT POLQ
▯ POLQ is able to bypass and extend cyclobutane pyrimidine dimers, but prefers purine over pyrimidine incorporation
▯ POLQ variants can protect against UV induced cell death.

## 1.1 INTRODUCTION

The genome is under perpetual bombardment by external and internal factors (i.e. electromagnetic radiation, environmental agents or metabolic components, respectively) that damage DNA. This damage fosters genomic instability, such as disruption of normal cell function through mutations and generation of single and double DNA strand breaks, ultimately leading to disease (such as cancer) or the triggering of cell death through apoptosis. While it is impossible for the cell to avoid DNA damage, maintaining a functional genome is imperative to the cell survival and function [1–3]. To combat the detrimental effects of DNA damage the cell employs a system of specialized DNA repair enzymes to fix damaged DNA and maintain a properly functioning genome.

The DNA repair mechanisms within the cell are composed of a series DNA polymerases that are specialized to repair the many types of DNA damage the cell encounters [4]. The combined activities of these repair mechanisms are highly effective and efficient. However, some mechanisms are more faithful at returning DNA to its original sequence (i.e. homologous recombination, HR) than others (i.e. nonhomologous end joining, NHEJ), which often generate mutations [5–7]. Despite these mutagenic behaviors, low fidelity mechanisms of DNA repair are beneficial to the cell as they repair broken DNA strands, often helping the cell avoid apoptosis due to DNA fragmentation [7,8].

Given the inherent nature of DNA repair mechanisms to promote cell survival while also promoting mutagenesis, albeit at a low level, factors involved with DNA repair pathways have long been linked to carcinogenesis [4,9]. To date, every known DNA repair polymerase, as well as many of their cofactors have been identified in cancer [10–15]. It is hypothesized that the ability of DNA repair mechanisms to help the cell avoid apoptosis while also generating genomic mutations drives carcinogenesis aiding in disease. This makes understanding how DNA repair polymerases work of great interest to understand their role in carcinogenesis. One such DNA repair enzyme that appears to be an emerging factor in melanoma is DNA polymerase theta (Pol θ or POLQ). Recently, variant forms of Pol θ were identified in a number of patient-derived melanoma samples [16]. An A-family DNA repair enzyme, Pol θ is inherently error prone and is essential for cell function and organismal development [6,7,17]. Pol θ plays a predominant role in two repair pathways, microhomology-mediated end joining (MMEJ, also known as theta-mediated end joining, (TMEJ), and translesion nucleotide bypass [18–21].

TMEJ is a highly error prone DNA double strand break (DSB) repair pathway that is activated when the predominant DSB repair pathway, homologous recombination (HR), is overwhelmed (such as when the genome occurs many DSBs) and/or inactive (such as in cancer states) [22–25]. Unlike in HR, a high-fidelity repair pathway, which uses the sister chromatid as a template for repair, TMEJ, uses small domains of microhomology between the two strands around the site of the DSB as sites of recognition for the repair machinery. Pol θ then generates a varying number of bases to repair the break. In the absence of a template, much of the new sequence represents mutations to the original genomic information. However, repairing the DSB is essential to genomic integrity and aids in avoiding DSB induced apoptosis. Additionally, Pol θ also functions in translesion nucleotide bypass, aiding stalled replisomes in circumventing bulky nucleotide lesions (i.e. cyclobutane pyrimidine dimers (CPD), 8-oxoguanine (8-oxoG), abasic (AP) sites) encountered during replication [8,21]. This is an essential function during replication. By allowing replicative DNA polymerases to continue replication, replication fork collapse and the subsequent DNA DSB is avoided. While bypassing the DNA damage without repair perpetuates genomic mutations, this avoids potentially greater mutagenesis due to TMEJ repair [26]. Together this indicates that Pol θ function is intrinsically mutagenic yet required for cell function. This duality, mutagenic enzymatic behavior while also supporting cell survival [17,27,28], along with aberrant Pol θ activity in cancer cells [29–31] has led many to hypothesize that Pol θ activity supports carcinogenesis. However, much of our information regarding Pol θ function comes from over- and loss-of-function studies, with very few determining how variant forms of Pol θ, like those found in melanoma tumors, may contribute to carcinogenesis.

It has been shown previously that patient derived variants display aberrant DNA polymerase activity when compared to that of wild-type Pol θ *in vitro* [32]. This included altered polymerization rates and decreased fidelity. Based on these differences it can be hypothesized that these variants perpetuate a disease state a promote cancer. However, as of yet there have not been any studies determining how these variants function within the cell or how or if the variants can bypass DNA damage such as CPD lesions which can be generated from exposure to UV.

Here we present our findings using variant forms of Pol θ from human melanoma samples to determine how variant Pol θ may contribute to carcinogenesis. We show that like wild-type Pol θ, variant forms of Pol θ can also perform translesion nucleotide bypass on CPD damaged DNA. We also show that overexpression of variant Pol θ rescues UV induced apoptosis.

## 1.2 MATERIALS AND METHODS

All materials were purchased from Sigma-Aldrich (St. Louis, MO), Bio-Rad Laboratories (Hercules, CA), AmericanBio (Canton, MA), and Research Products (Mount Prospect, IL). DNA oligonucleotides were purchased from Integrated DNA Technologies (Newark, NJ) and deoxynucleotides from New England Biolabs (Ipswich, MA). All DNA oligos were purified via HPLC with standard desalting from the manufacturer. Cell lines were obtained through ATCC.

### 1.2.1 Purified Protein and DNA Substrates

Recombinant pSUMO3 plasmids containing either WT or mutated the truncated polymerase domain were expressed in *E. coli* and purified as previously described [32,33]. The double-stranded DNA substrates (dsDNA) were generated by annealing complementary deoxynucleotide oligos purchased from IDT. Substrates containing cyclobutane pyrimidine thymine-thymine dimers (CPD) representing damaged DNA, were synthesized in the Delaney laboratory (Sarah Delaney, Brown University) as previously described [33].

The 5’6-FAM primers were annealed to complementary DNA templates with sequence context as previously described [7,34] and are described below:

24/33 undamaged DNA substrate

5’-/FAM/-TTT CTC CGG TAC TCC AGT GTA GGC GAG GCC ATG AGG TCA CAT CCG TTA GCT TGA GAA

24/33 CPD damaged DNA substrate where the underlined <u>TT</u> indicates the CPD lesion 5’-/FAM/-TTT CTC CGG TAC TCC AGT GTA GGC GAG GCC ATG AGG TCA CAT CCG <u>TT</u>A GCT TGA GAA

Confirmation of annealed substrates was determined with 12% Native PAGE scanned on an RB Amersham Typhoon Fluorescent Imager (Cytiva) with a FAM filter.

### 1.2.2 Electromobility shift assay

The DNA binding affinity constant K_D(DNA)_ was determined for the variants tested as previously described [35]. Pol θ variants were titrated from 0-1800 nM against 10nM 25/40 dsDNA substrate in binding buffer (10 mM Tris pH 7.0, 6 mM MgCl_2_,100 mM NaCl, 10% Glycerol, 0.1% NP40) and incubated for 1 hour at room temperature. Samples were separated on a 6% Native PAGE and scanned on an RB Typhoon scanner (Cytiva) with the FAM fluorescence filter. Separated bound and unbound products were quantified using ImageQuant. K_D(DNA)_ was determined by equation 1.

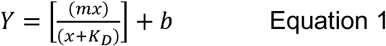

At minimum, three replicates and two protein preparations were used to generate this data.

### 1.2.3 Primer extension assay

Qualitative primer extension assays were performed as previously described [32,35]. Briefly, Pol θ (750 nM) was pre-incubated with 50 nM CPD damaged DNA and incubated for 5 minutes at 37°C (Pol-θ/DNA). MgCl_2_ (10 mM), 50 μM dNTP (either no nucleotides (none), all, or individual) were added to Pol θ/DNA reaction mixture in buffer containing 20 mM Tris HCl, pH 8.0, 25 mM KCl, 4% glycerol, 1 mM βME, and 80 μg/mL BSA to initiate the reaction and incubated at 37°C for up to 5 minutes and quenched with 80% formamide/EDTA. Products were separated out on a 15% urea-denaturing PAGE and scanned on an RB Amersham Typhoon fluorescent imager (Cytiva). Single turnover experiments were carried out in a similar manner except with 50 nM CPD damaged DNA substrate and 200 nM Pol θ. Nucleotides were titrated from 0-5000 μM. Pol θ/CPD DNA substrate and nucleotide mixtures were preincubated separately for 5 mins at 37°C. Once combined, samples were incubated at various times for up to 5 mins at 37°C and reactions stopped with 80% formamide/100 mM EDTA. Products were separated and visualized as described above. Data was fit to a single exponential equation (Equation 2) to determine *k_obs_* for each concentration. The rate was then graphed against varying nucleotide concentrations to determine *k_pol_* and K_d(dNTP)_ as previously described (Equation 3) [32]. Experiments were repeated 3-5 times with a minimum of two protein preparations by two different individuals to determine rates (± standard error).

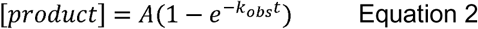

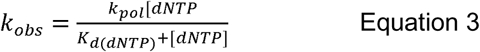

### 1.2.4 Western blot analysis

Briefly, protein samples were loaded into polyacrylamide gels and subjected to sodium dodecyl sulphate–polyacrylamide gel electrophoresis. To detect the protein of interest, the membranes were incubated with antibodies recognizing POLQ (custom below, Pacific Immunology) and α-Tubulin (Cell Signaling Technology 2144S). The secondary antibodies used were anti-rabbit IgG, HRP-linked antibody (Sigma A4914) and anti-mouse IgG, HRP-linked antibody (Sigma A9917).

### 1.2.5 Generation of POLQ antibody

For this study a monospecific POLQ antibody was generated by Pacific Immunology in rabbits using a synthesized peptide for the POLQ polymerase domain: CSIFRARKRASLDINKEKPG. Antibody was affinity purified from whole serum.

### 1.2.5 Survival experiments

siRNA oligos targeting POLQ 3’ UTR (one uniquely designed by our group (the first one) a second from previously published work [36] (the second) and a scrambled control were obtained from Dharmacon (Horizondiscovery.com).

siRNA sequences:

1. 5’ GGGUAAACCAUGAAGAAAA 3’
2. 5’ GAGAUUACCCUUUCACCUA 3’

Western blot analysis for POLQ knock down was performed and based on these results, the 3’ UTR siRNA was used for our experiments indicating a partial knock down and rescue of POLQ (sfig 1).

Cell survival was measured using the Cell TiterGlo assay (Promega) following the manufacturers protocol. MCF7 (ATCC HTB-22) or dermal melanocyte (ATCC CRL-4059) cells were dispensed into 96-well plates and grown to 80-90% confluency. Cells were then co-transfected with 50-100 nM of siRNA directed to POLQ or a scrambled control and 2 μg of a POLQ rescue plasmid using nucleofection for the MCF7s and lipofectamine 2000 for the dermal melanocytes, following manufacturer’s instructions. WT POLQ rescue plasmids contained full-length WT POLQ obtained from Addgene (Addgene plasmid # 73132; http://n2t.net/addgene:73132; RRID:Addgene_73132, [37]). Variant POLQ rescue plasmids were generated using Site Directed Mutagenesis as previously reported [32]. Transfected cells were incubated for 48-72 hours at 37 °C. Media was then replaced with PBS and cells were exposed to either 5, 10, or 20 J/m^2^ UV. After UV exposure the PBS was replaced with complete media and cells were allowed to recover for 24 hours. Cells were then assayed for adenosine triphosphate luminescence using Cell TiterGlo reagent as described in the manufacturer’s instructions using a Spectra Max Gemini XPS microplate reader from Molecular Devices. 5 J/m^2^ UV exposure produced a repeatable 50% cell survival of POLQ siRNA knock-down cells and was used as our exposure intensity for the experiments.

### 1.2.7 Comet Assay

MCF7 cells were co-transfected with 50-100 nM siRNA for POLQ and 2 μg of rescue plasmids for either WT POLQ, T2161I, E2406K, or L2538R using nucleofection incubated for 48-72 hours. For UV exposure, media was replaced with PBS and cells were exposed to 5 J/m^2^ UV. Following UV exposure, PBS was replaced with complete media and cells were incubated for 24 hours. The cells were prepared and analyzed in a neutral comet assays, according to published procedures [38] using Comet slides (Trevigen Cat # 4250–200-03). Image analysis of at least 50 cells was performed using CometScore software (TriTek, Sumerduck, VA, USA). Data are represented as mean ± SD (n = 50).

## 1.3 RESULTS

### 1.3.1 DNA polymerase theta cancer variants express a strong affinity for CPD damaged DNA

The first step in the DNA polymerization pathway is to bind DNA [32]. To assess the ability of Pol θ to bind to either undamaged or CPD damaged DNA, we performed an electrophoretic mobility shift assay (EMSA) in which we titrated Pol θ against 5’ FAM labeled DNA substrate and observed DNA-Pol θ complexes. An apparent equilibrium dissociation constant (K_D(DNA)_) was determined by observing the concentration of Pol θ in which the complex was half bound. WT Pol θ bound undamaged DNA at 94 nM, whereas the variants experienced tighter binding of undamaged DNA at around 50 nM. In the presence of CPD damaged DNA, WT Pol θ experienced a five-fold increase in affinity (Fig 1, Table 1). Although L2538R, E2406K, and T2161I, had a slight reduction in affinity for undamaged DNA; the affinity increased significantly (from 11-36 fold) for CPD damaged DNA indicating much tighter binding compared to WT Pol θ.

**Figure One.**
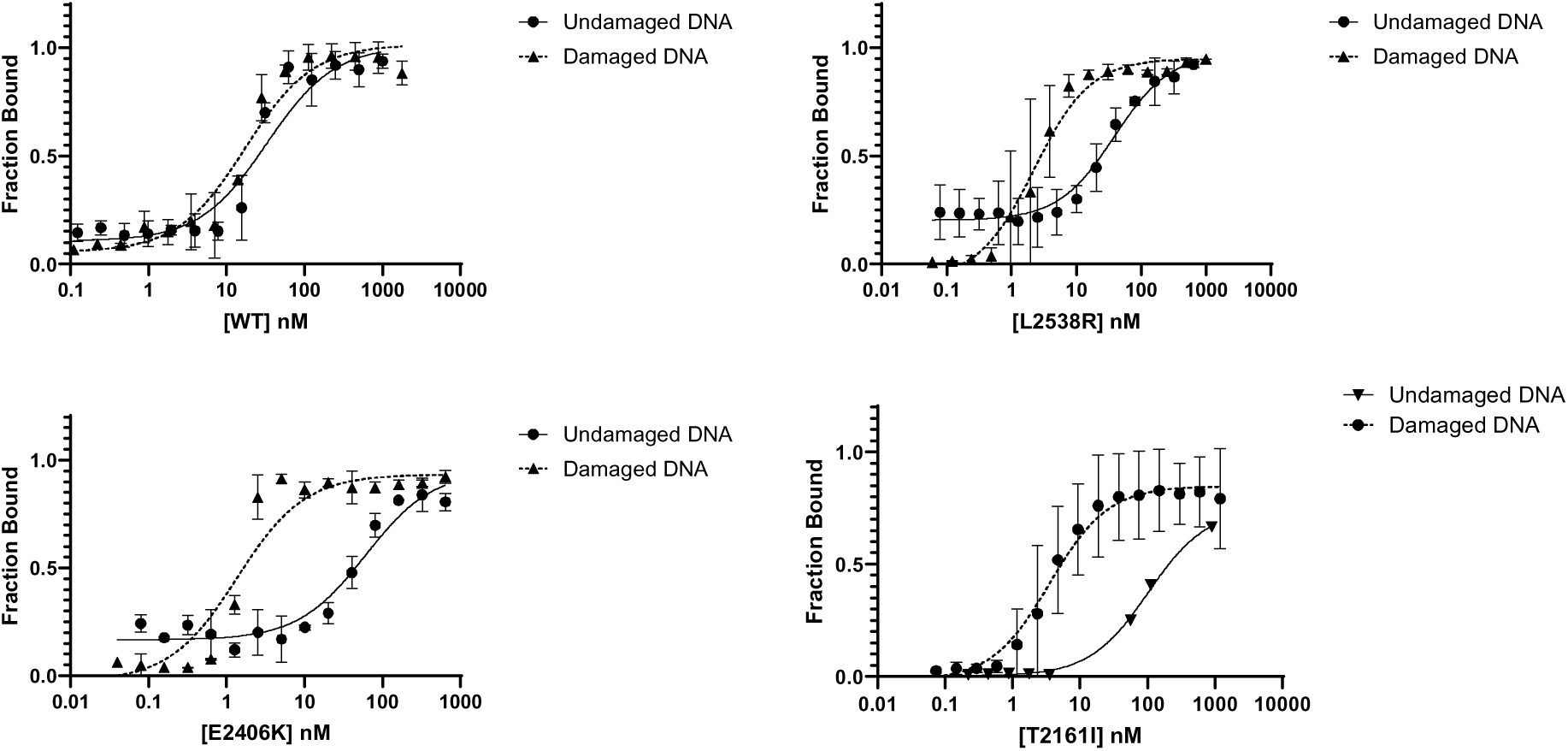
Variants display higher affinity for CPD damaged DNA compared to WT Pol θ. WT POLQ and L2538R, E2406K, and T2161I variants were titrated from 0-1800 nM against 10 nM undamaged (circles) or damaged (triangles) dsDNA. Bound and unbound products were separated on a 6% non-denaturing gel and quantified using ImageQuant software. K_D(DNA)_ was calculated using equation 1 (Methods).

**Table 1.** POLQ affinity for damaged and CPD damaged DNA.

| | Undamaged | Damaged | $\Delta$ Fold Change<sup>a</sup> |
| --- | --- | --- | --- |
| WT | 93.9 ± 14.7 nM | 17.3 ± 3.3 nM | 5 |
| L2538R | 49.8 ± 11.8 nM | 2.4 ± 0.6 nM | 21 |
| E2406K | 51.0 ± 10.8 nM | 1.4 ± 0.3 nM | 36 |
| T2161I | 48.2 ± 8.3 nM | 4.4 ± 1.1 nM | 11 |

<sup>a</sup> Fold change = undamaged/damaged

### 1.3.2 Cancer associated variants are able to bypass and extend CPD damaged DNA

To assess the ability of WT Pol θ and the cancer associated variants to bypass and extend CPD damaged DNA we incubated Pol θ with the 5’-FAM labeled CPD damaged DNA under single turnover conditions (excess enzyme) with either all of the nucleotides or the individual nucleotides for five minutes at 37°C. When given all nucleotides both WT Pol θ and the cancer associated variants incorporated dATP opposite the CPD lesion and then extend to the full-length product of n+12 (Fig 2). With dATP only, WT Pol θ and T2161I were able bypass and extend, whereas L2538R and E2406K were able to bypass and display minimal extension. No bypass was observed for any Pol θ (WT and variants) with dCTP. For dTTP, Pol θ (WT and variants) only incorporated opposite the first T of cyclobutane pyrimidine dimer (n+1). For dGTP, extension was only observed with the T2161I variant, with minimal extension observed with WT Pol θ, L2538R, and E2406K.

**Figure Two.**
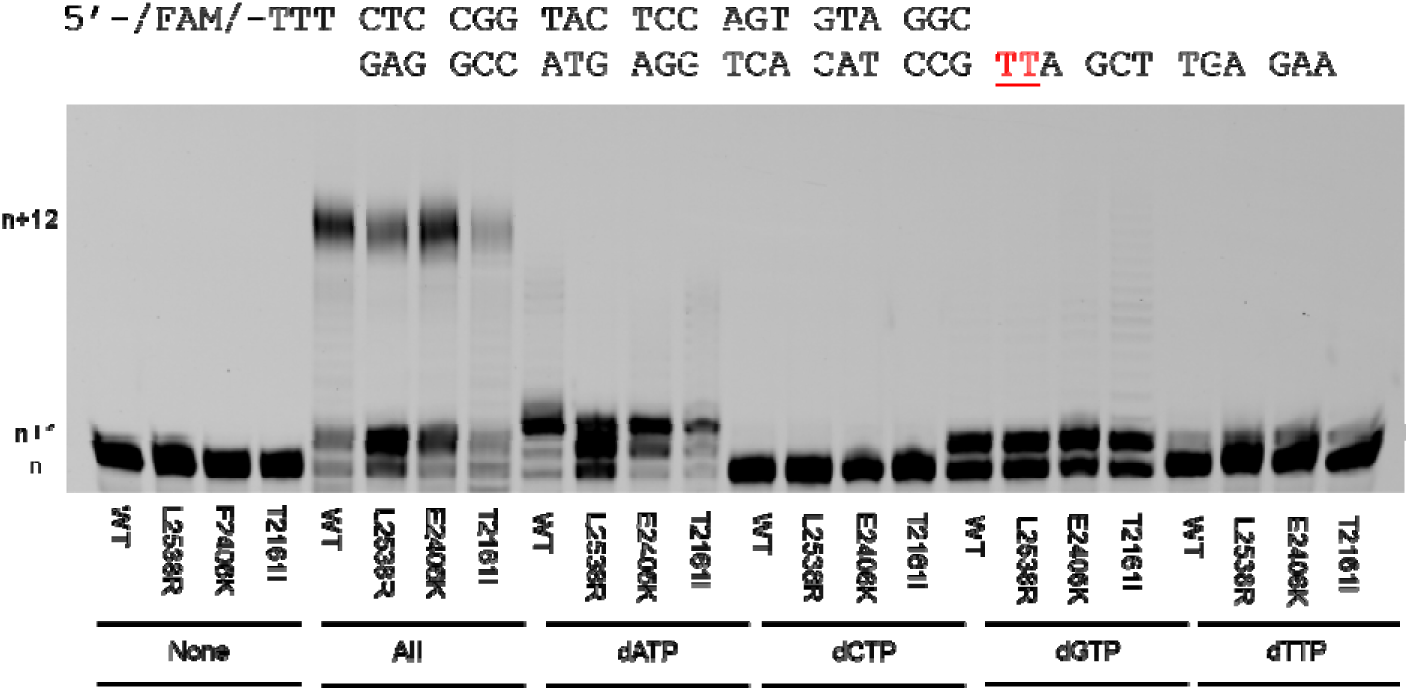
WT Pol θ and cancer variants bypass and extend past CPD damaged DNA. A representative denaturing gel demonstrates primer extension of CPD dsDNA. Under single-turnover conditions 750 nM of WT POLQ and L2538R, E2406K, and T2161I variants were preincubated with 50 nM of CPD damaged dsDNA substrate and then combined with 10 mM MgCl_2_ and either no dNTPs or 50 µM of All dNTPs, dATP, dCTP, dGTP, or dTTP for 5 minutes at 37°C. DNA extension products were separated on a denaturing gel and visualized on a Typhoon scanner. Each n+1 band represents extension past the CPD damage with either correct (dATP) or incorrect incorporation (dCTP, dGTP, or dTTP). Each additional band represents another extension with a maximum product of n+12.

### 1.3.3 Cancer associated variant L2538R displays higher fidelity during CPD bypass but is less efficient compared to WT Pol θ

While primer extension assays qualitatively define polymerase activity, we investigated time-based single-turnover kinetics to determine the polymerization rate (*k_pol_*) and apparent equilibrium dissociation constant (K_d(dNTP)_) during bypass of the CPD lesion. The reaction conditions contained a 4:1 ratio (protein:DNA) with excess enzyme while titrating individual nucleotide concentration. Pol θ/CPD damaged DNA were preincubated and were mixed with varying concentrations of nucleotide for 0-5 minutes at 37°C. Products were separated on denaturing gel and quantified as described (methods).

Data from our single turnover experiments are summarized in Table 2. WT Pol θ incorporation of dATP (correct) opposite the TT dimer had a kpol of 0.13 s^-1^ with a K_d(dNTP)_ of 26 μM. For dGTP (incorrect) opposite the TT dimer the rate remained the same as correct, but the nucleotide affinity decreased by 4-fold. For dTTP (incorrect) the rate decreased to 0.03 s^-1^ and the K_d(dnTP)_ value increased to 143 μM (a 5 fold decrease in affinity). Interestingly, we were unable to quantify any data using dCTP (incorrect). This data is consistent with the qualitative primer extension assays (Fig 2).

**Table 2.** Single turnover kinetics for POLQ with CPD damaged DNA^a^.

| Protein | Sequence | $k_{pol} (\text{s}^{-1})$ | $K_d (\mu\text{M})$ | Efficiency<sup>b</sup><br>$k_{pol}/K_d (\mu\text{M}^{-1} \text{s}^{-1})$ | Δ-fold for<br>efficiency<sup>c</sup> | $F^d$ | Δ-fold<br>fidelity<sup>e</sup> |
| --- | --- | --- | --- | --- | --- | --- | --- |
| hWT | TT:dATP | $0.13 \pm 0.005$ | $25.7 \pm 5.6$ | 0.0050 | | | |
| hWT | TT:dGTP | $0.13 \pm 0.005$ | $98.1 \pm 21.6$ | 0.0013 | | 4.9 | |
| hWT | TT:dTTP | $0.04 \pm 0.004$ | $143.2 \pm 22.6$ | 0.0002 | | 7.1 | |
| hWT | TT:dCTP | Undefined | undefined | -- |  | -- |  |
| L2538R | TT:dATP | 0.14 ± 0.005 | 80.1 ± 12.0 | 0.0017 | 3 |  |  |
| L2538R | TT:dGTP | 0.04 ± 0.002 | 198.8 ± 43.3 | 0.0002 | 7 | 10.1 | 0.5 |
| L2538R | TT:dTTP | 0.02 ± 0.001 | 496.3 ± 84.0 | 0.00004 | 5 | 43.7 | 0.2 |
| L2538R | TT:dCTP | Undefined | undefined |  |  |  |  |
| E2406K | TT:dATP | 0.18 ± 0.006 | 47.4 ± 8.6 | 0.0037 | 1 |  |  |
| E2406K | TT:dGTP | 0.11 ± 0.004 | 79.1 ± 15.7 | 0.0014 | 1 | 3.6 | 1.4 |
| E2406K | TT:dTTP | 0.05 ± 0.008 | 1144 ± 143 | 0.00004 | 114 | 86.4 | 0.1 |
| E2406K | TT:dCTP | Undefined |  |  |  |  |  |
| T2161I | TT:dATP | 0.31 ± 0.012 | 102.1 ± 13.2 | 0.0031 | 1.6 |  |  |
| T2161I | TT:dGTP | 0.14 ± 0.008 | 259.4 ± 62.8 | 0.0006 | 2.3 | 6.5 | 0.8 |
| T2161I | TT:dTTP | Undefined | undefined |  |  |  |  |
| T2161I | TT:dCTP | Undefined | undefined |  |  |  |  |

<sup>a</sup>Polymerization rates and dissociation constants derived from single-turnover experiments for WT and cancer-associated variants (± standard error).
<sup>b</sup> $k_{pol}/K_d$ ( $\mu\text{M}^{-1}\text{s}^{-1}$ ).
<sup>c</sup> $\text{efficiency}_{WT}/\text{efficiency}_{variant}$ .
<sup>d</sup> $F = (\text{efficiency}_{correct} + \text{efficiency}_{incorrect})/\text{efficiency}_{incorrect}$ .
<sup>e</sup> $\text{fidelity}_{WT}/\text{fidelity}_{variant}$ .

The *k_pol_* rates for L2538R and E2406K were similar to WT Pol θ with correct nucleotide incorporation but we observed a 3-fold weaker affinity for correct nucleotide with L2538R and a 2-fold reduction in K_d(dNTP)_ for E2406K compared to WT Pol θ. For incorrect nucleotide dGTP, we observed both a reduced polymerization rate and nucleotide affinity for L2538R whereas E2406K incorporation rate was similar to WT Pol θ. Like WT Pol θ, no values were obtained for dCTP for L2538R and E2406K. Incorporation of dTTP was slow for WT, L2538R and E2406K, but the affinity was greatly reduced with up to 3-fold less affinity for dTTP for L2538R and 8-fold reduction for E2406K. Comparing all these values to WT Pol θ, we observed that L2538R is less efficient (3-7 fold) at incorporating nucleotides but experiences greater fidelity when bypassing CPD lesions whereas E2406K displayed a reduction in efficiency with dTTP. T2161I incorporated dATP (correct) at almost double the *k_pol_* rate and dGTP incorporation at a similar polymerization rate compared to WT Pol θ but again, with a weaker affinity for nucleotides. Quantification of T2161I dTTP incorporation was inconclusive resulting in an unstable mathematical fit of the data and we observed no product formation with dCTP. Like E2406K, T2161I experienced similar fidelity and efficiency compared to WT Pol θ.

### 1.3.4 Pol θ variants partially rescue UV induced apoptosis in Pol θ RNAi treated cells

One key role of Pol θ in the cell is to protect the cell from DSB induced apoptosis while homologous recombination (the favored pathway) is overwhelmed [39]. Previous observations have shown that in the absence of Pol θ, cells are more sensitive to UV radiation, undergoing apoptosis at a higher rate than cells with Pol θ [7]. To determine whether the variants were still able to protect against UV radiation we performed RNAi induced knockdown of endogenous Pol θ and rescued Pol θ with either full-length wildtype or cancer derived Pol θ variants. Cells were co-transfected with either siRNA oligos to the 3’ UTR of Pol θ or a scrambled control oligo and either a WT or variant rescue plasmid encoding full-length Pol θ. For these experiments we used two types of stable transformed cells, dermal melanocytes (DM) and MCF7 cells, a breast cancer cell line.

Upon UV exposure in cells transfected with only the scrambled siRNA oligo we observed 85% and 80% cell survival in the MCF7 and DM cells (respectively, Fig 3). In these cells when transfected with siRNA targeting Pol θ alone and exposed to UV, we observed a further drop in cell survival, with survival rates of 55% and 50% (MCF7 and DM respectively). In MCF7 and DM cells co-transfected with the Pol θ siRNA oligo and the WT Pol θ rescue construct we observed a significant increase in cell survival compared to cells transfected with the Pol θ siRNA alone, with nearly 75% and 80% survival (respectively).

**Figure Three.**
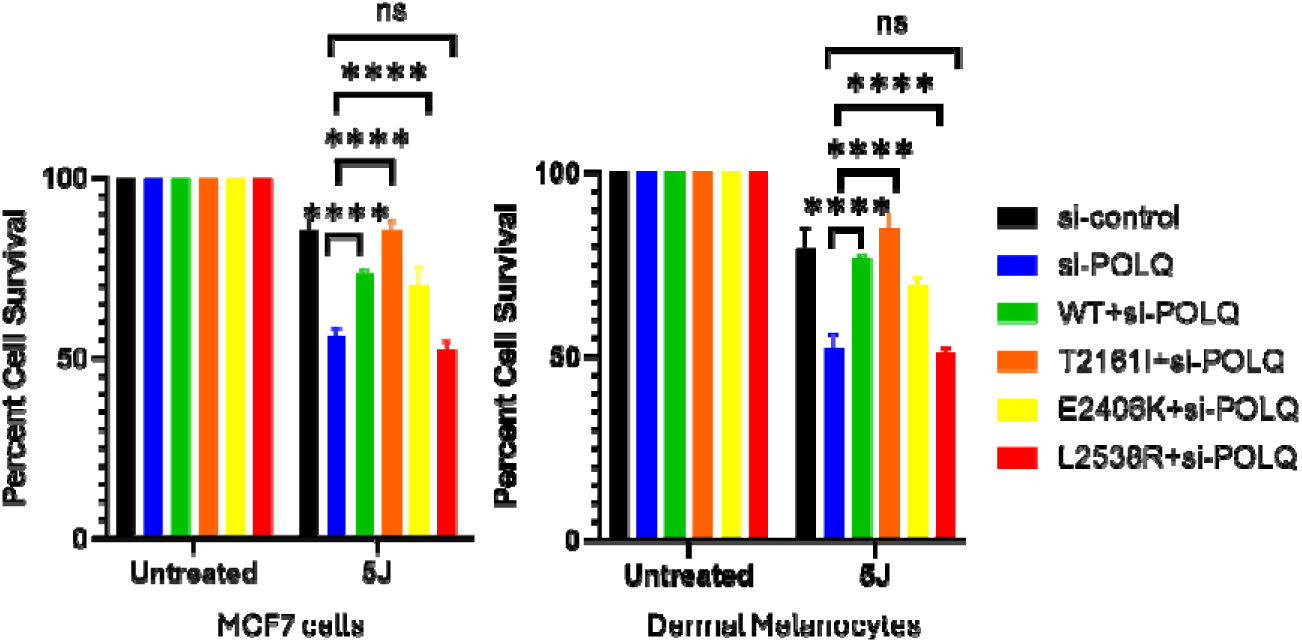
Pol θ variants rescue UV induced cell death. MCF7 and dermal. Cells were harvested using Cell TiterGlo and adenosine triphosphate luminescence was quantified using a microplate reader. Significance was determined using Two-way ANOVA (**** p < 0.0001).

We next compared the ability of Pol θ variants T2161I, E2406K and L2538R to rescue the loss of cell viability observed in cell transfected with Pol θ siRNA and exposed to UV radiation. Using rescue constructs containing full-length variant forms of Pol θ co-transfected with the Pol θ siRNA oligos we were able to observe significant rescue for cells transfected with T2161I (85% survivability for both MCF7 and DM cells) and E2406K (nearly 75% and 70% survivability MCF7 and DM cells respectively). In both MCF7 and DM cells co-transfected with siRNA and L2538R, survivability was not significantly changed when compared to the Pol θ siRNA alone. These results indicate that T2161I and E2406K variants were still able to protect cells from UV induced cell death, similar to that of WT Pol θ, while L2538R could not.

To better understand why L2538R was unable to rescue the cell survival phenotype like the other variants we performed a comet assay to quantify the number of DSBs in the UV treated cells. Though UV radiation does not directly cause DNA DSBs it is hypothesized that a large amount of DNA damage could overwhelm DNA repair pathways, leading to secondary DNA damage, such as DSBs due to replication fork collapse [40,41]. In this case replication fork collapse presumably occurs due to failure to bypass CPD lesions which stall replicative polymerases.

Similar to the rescue experiments, MCF7 cells were either exposed to UV radiation or not after being transfected with either a scrambled siRNA control or Pol θ siRNA oligos as well as either no Pol θ rescue construct, a full-length WT Pol θ rescue construct, or one of the variant full-length Pol θ rescue constructs (Fig 4). In cell transfected with the scrambled siRNA oligo and exposed to UV radiation, we observed no significant difference in the number of double strand breaks compared to cells not treated with UV radiation. Cells transfected with the Pol θ siRNA oligo and treated with UV displayed a significant increase in observable DSBs when compared to the control siRNA transfected cells. This was rescued however in cells co-transfected with the Pol θ siRNA oligo and the WT Pol θ rescue construct, which had no significant increase in DSBs when compared to the UV exposed and scrambled siRNA cells.

**Figure Four.**
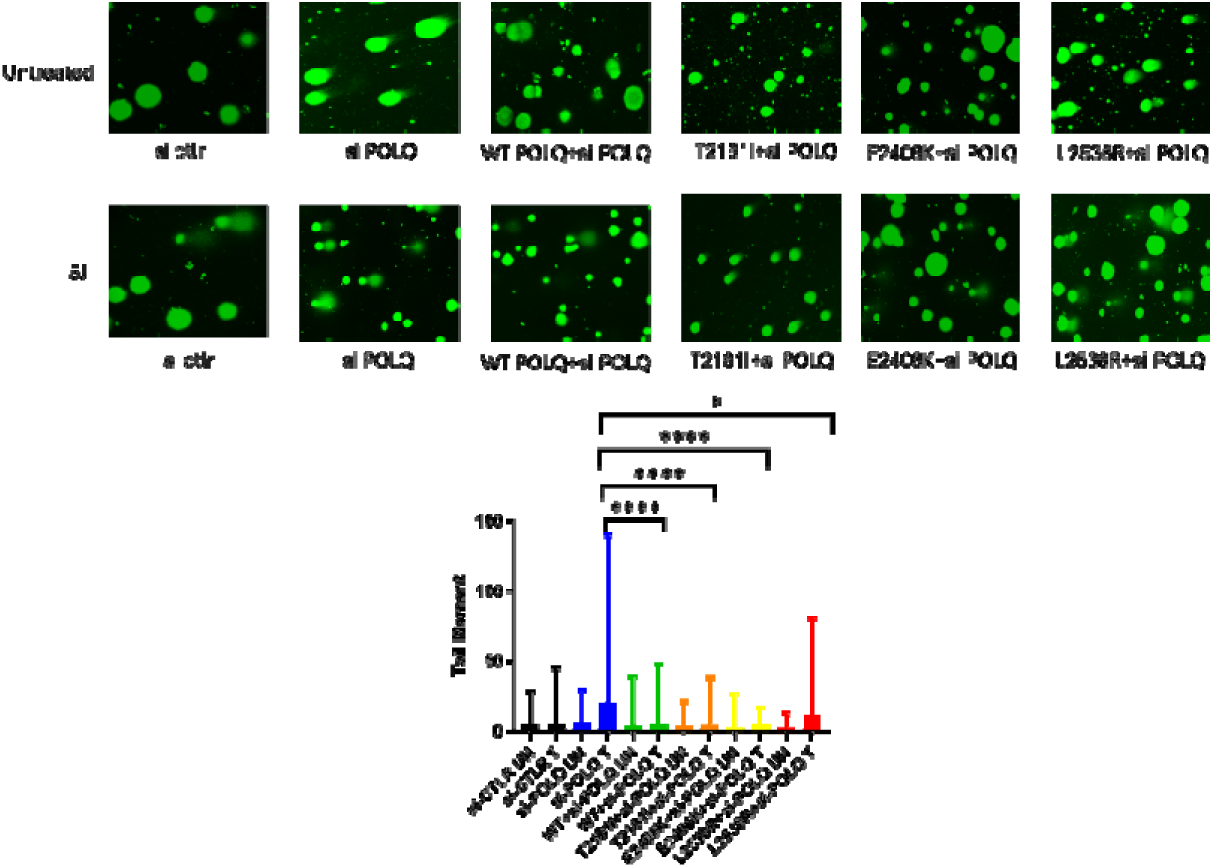
WT POLQ and variants E2406K, and T2161I reduce the number of double strand brakes in UV treated cells. MCF7 cells melanocytes were co-transfected with either a scrambled control siRNA oligo or an siRNA oligo to POLQ and either a siRNA resistant WT, L2538R, E2406K, or T2161I POLQ rescue plasmid. Cells were allowed to grow for 48-72 hrs. and then treated with either 5J ultra-violet radiation or not and allowed to recover for 24 hrs. Neutral comet assays were performed using Comet slides (n ≥ 50 cells) and analyzed using CometScore software. Error bars represent standard deviation, and significance was determined using One-way ANOVA (**** p < 0.0001).

We next compared these results to cells co-transfected with the Pol θ variant rescue constructs and Pol θ siRNA oligo. Compared to the scrambled siRNA oligo treated with UV, we saw no observable change in DSB formation in cells treated with UV and co-transfected with Pol θ siRNA and either T2161I or E2406K variant rescue constructs. However, we did observe a significant increase in the number of DSBs observed in UV treated cell that were co-transfected with Pol θ siRNA and the L2538R variant repair construct. These data suggest that WT Pol θ and variants T2161I and E2406K aid in genomic stability by reducing the number of DSBS breaks in the cell. In addition, though these data are not conclusive, these results indicate that L2538R containing cells have a larger number of DSBs in the absence of WT Pol θ function when treated with UV radiation.

## 1.4 DISCUSSION

Understanding the biochemical behavior of a DNA polymerase can give valuable insight into the function of the DNA polymerase in the cell. In that same way, defining the biochemical characteristics of aberrant DNA polymerase variants identified in cancer can provide valuable information to better understand carcinogenesis. For our study we used biochemical and cell-based techniques to characterize how Pol θ variants, L2538R, E2406K, and T2161I, identified from patient melanoma tumors [16,32] behaved during DNA repair in the presence of CPD lesions and UV damaged DNA.

Previous work has shown that the three variants tested here displayed aberrant polymerase characteristics compared to WT Pol θ in the presence of undamaged DNA [32]. Though these studies provide valuable information regarding the behavior of variant forms of Pol θ, it does not address how these variants would behave in cell when confronted with damaged DNA. Our work here provides insights into how these variant forms may contribute to genomic stability and cancer progression.

### 1.4.1 Cancer variants have a higher affinity for damaged DNA that WT Pol θ

To our knowledge no study has looked at the capabilities of variant forms of Pol θ to repair damaged DNA. The first step of DNA repair for Pol θ is binding damaged DNA. Using EMSAs we determined that the cancer associated variants displayed a significantly higher affinity for damaged DNA compared to WT Pol θ (Fig 1, Table 1). These data indicate that the cancer variants in addition to binding undamaged DNA (as previously identified) could still bind to damaged DNA. Perhaps most surprising is that this increased affinity for damaged DNA is so great, particularly given the nature of the mutations (all missense) and that all the mutations come from different regions of the polymerase domain (L2538R = palm, E2406K = fingers, and T2161I = thumb) not just the thumb domain which is associated with DNA binding.

### 1.4.2 Cancer variants bypass and extend CPD damaged DNA with L2538R displaying greater fidelity but lower efficiency

Previous studies have shown that Pol θ can bypass some DNA lesions including 8-oxoguanine, apurinic sites, thymine glycol, and limited activity on Cisplatin damaged DNA [33,42,43], but is unable to bypass photoadducts such as CPD and 6-4 photoproduct (6,4 PP) under steady-state (excess DNA substrate) conditions [43]. A further study looked at the ability of Pol θ to bypass and extend past CPD and 6-4 PP lesions (also under steady state conditions) with the help of Pol ι or in the presence of a primer that overlapped the damage by one base, suggesting that Pol θ could not initiate bypass over a photoadduct lesion let alone extend past the damage [21,44]. Here however, we observed that Pol θ as well as the variants, under single-turnover conditions, are able to initiate bypass across CPD lesions (n+1) as well as extend past it to generate full-length product (n+12) (Fig 2). By carrying out nucleotide incorporation experiments under single-turnover conditions, we can directly observe the turnover of the substrate to n+1 without complications of steady-state product accumulation that would otherwise obscure this product formation [32,45].

Moreover, we utilized single-turnover conditions further to better understand the mutagenic tendencies of Pol θ and its variants using time-based primer extension assays to understand the rate and nucleotide preference of Pol θ. The hypothesis is that Pol θ and/or its variants contribute to genomic instability by inserting any nucleotide across the photoadduct. However, our observations (Table 2) suggest that Pol θ and the variants tested here prefer purines when bypassing CPD damage. Additionally, neither WT Pol θ nor the cancer variants were able to incorporate dCTP and had such a low affinity for dTTP. This was interesting given our previous observations that Pol θ displayed little discrimination between purines and pyrimidines on undamaged substrate [32,33].

Though our data does not provide a reason why Pol θ cannot incorporate all nucleotides opposite the CPD lesion, we hypothesize the preference for purines follows standard Watson-Crick pairing to maximize hydrogen bonding stabilization and that the CPD:dCTP pairing instability was too great for binding and incorporation. Moreover, while E2406K, and T2161I showed similar rates for polymerization and nucleotide binding, variant L2538R demonstrated a significant decrease in efficiency. This suggests that the inherent inefficiency of L2538R to incorporate any nucleotide would contribute to genomic instability through replisome stalling and DNA DSB formation proceeded by replication fork collapse rather than mutagenesis.

### 1.4.3 Pol θ variant L2538R fails to rescue UV induced apoptosis

Prior to this work there have been no studies assessing the role of variant forms of Pol θ in the cell. Previous loss-of-function (LOF) studies have shown that in the absence of Pol θ cells are more sensitive to UV radiation [7]. To determine if the variants were able to confer the same level of protection we performed rescue experiments, partially knocking down endogenous Pol θ and over expressing a rescue Pol θ construct (Fig 3). Tested in two different cell lines, when endogenous Pol θ was partially knocked down and cells were exposed to UV radiation, we observed a reduction of cell survival, recapitulating the sensitivity previous observed in other Pol θ LOF studies. This sensitivity however was rescued when a full-length WT Pol θ construct was introduced, indicating that Pol θ function protects against UV induced apoptosis. When we preformed the same Pol θ LOF and rescued with the full-length Pol θ cancer variants, we observed rescue only with E2406K and T2161I. Cells supplied with the L2538R rescue construct were still sensitive to UV radiation, indicating that L2538R could not rescue cell death.

Based on our previous observation of L2538R we next wanted to determine if the cells containing L2538R were experiencing greater instances of DNA DSBs (Fig 4). Co-transfecting MCF7 cells with siRNAs to endogenous Pol θ and rescue plasmids for either WT Pol θ or one of the cancer variants. We observed that cell containing either WT Pol θ or variants E2406K or T2161I showed no significant increase in DNA DSBs when compared to the scrambled siRNA control when exposed to UV radiation. This indicated that variants E2406K and T2161I behaved like WT Pol θ supports our previous observations in the rescue experiments (Fig 3) suggesting that E2406K and T2161I can protect the cell against UV radiation. However, cells containing L2538R exposed to UV radiation had a significant increase in the number of DNA DSBs compared to the scrambled siRNA control. These observations support our hypothesis that L2538R cells have more DNA DSBs and though doesn’t explain why L2538R containing cells are unable to rescue UV induced cell death, supports our biochemical data that suggests L2538R’s decreased efficiency could lead to an increase in genomic stability.

In summary, the observations presented here suggest that though variants E2406K and T2161I have aberrant polymerase dynamics compared to WT [32] they still have overall function that is similar to WT Pol θ. This is particularly evident in their ability to protect the cell against UV radiation presumably through the reduction of DNA DSBs. However, this also implies that though these variants protect against cell death they are also promoting genomic instability in the form of genetic mutations through error prone repair. In a tumor of mixed cell populations and cells that contain both WT and variant Pol θ activity this suggests that cells with some variant forms of Pol θ could at least be as functional as cells with WT Pol θ and could potentially be more likely to progress disease due to their increased mutagenic capabilities.

The behavior of E2406K and T2161I contrasts with the observations for L2538R. Though L2538R is able to bind damaged DNA and bypass and extend like the other variants, it does so at a significantly reduced rate. Our data shows that L2538R cells have a greater number of DNA DSBs compared to WT Pol θ and the other variants tested, suggesting that this reduced rate contributes to replication fork collapse. This result though may not be very surprising given the location of the mutation in the palm domain of Pol θ, a key site for enzymatic activity within the protein. Future studies should aim at better understanding the role of Pol θ to promote genomic instability in the form of genetic mutations while also protecting against cell death through DSBs. Analysis of patient derived variants such as these have the potential to provide invaluable insight into functionally important mutations that promote cancer and aberrant function that could be exploited as makers and therapeutic targets.

## Notes

### Competing Interest Statement

The authors have declared no competing interest.

### Funding

- National Institute of General Medical Sciences

