## Supplemental data for "POLQ variants with aberrant DNA polymerase activity protect against UV-induced cell death"

### Slide 1
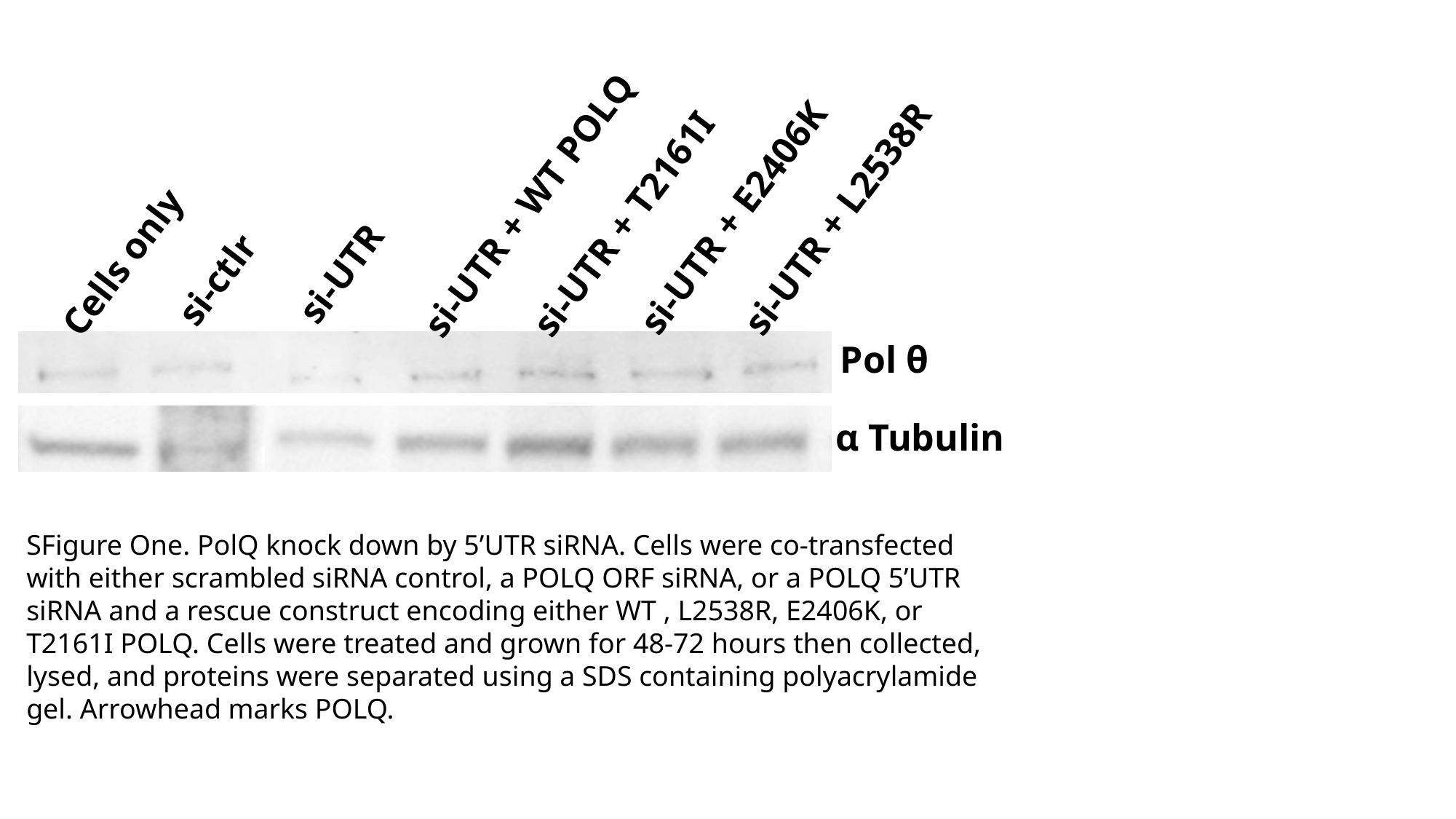

si-UTR + WT POLQ
si-UTR + E2406K
si-UTR + L2538R
si-UTR + T2161I
Cells only
si-UTR
si-ctlr
Pol θ
α Tubulin
SFigure One. PolQ knock down by 5’UTR siRNA. Cells were co-transfected with either scrambled siRNA control, a POLQ ORF siRNA, or a POLQ 5’UTR siRNA and a rescue construct encoding either WT , L2538R, E2406K, or T2161I POLQ. Cells were treated and grown for 48-72 hours then collected, lysed, and proteins were separated using a SDS containing polyacrylamide gel. Arrowhead marks POLQ.

### Slide 2
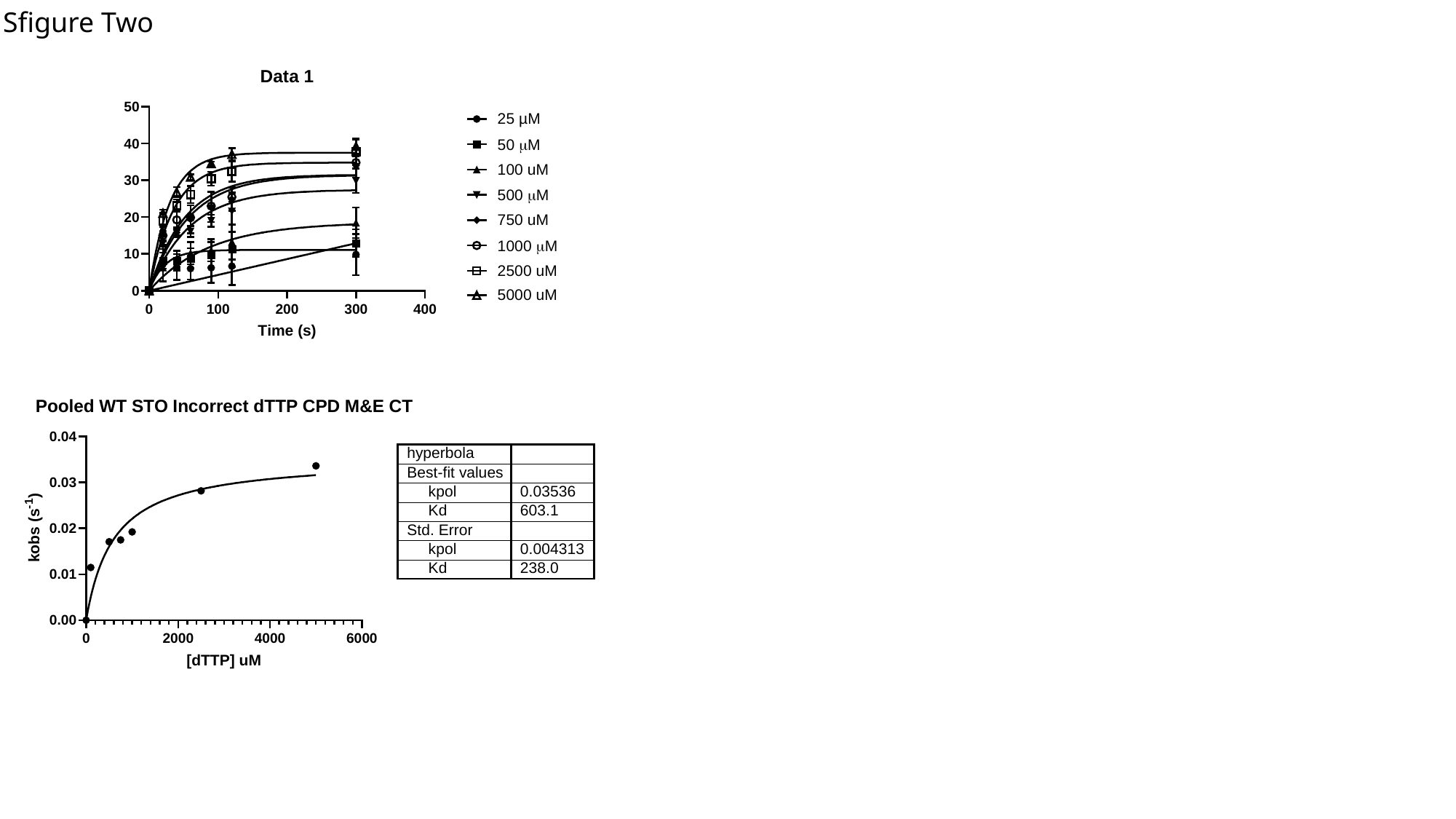

Sfigure Two

### Slide 3
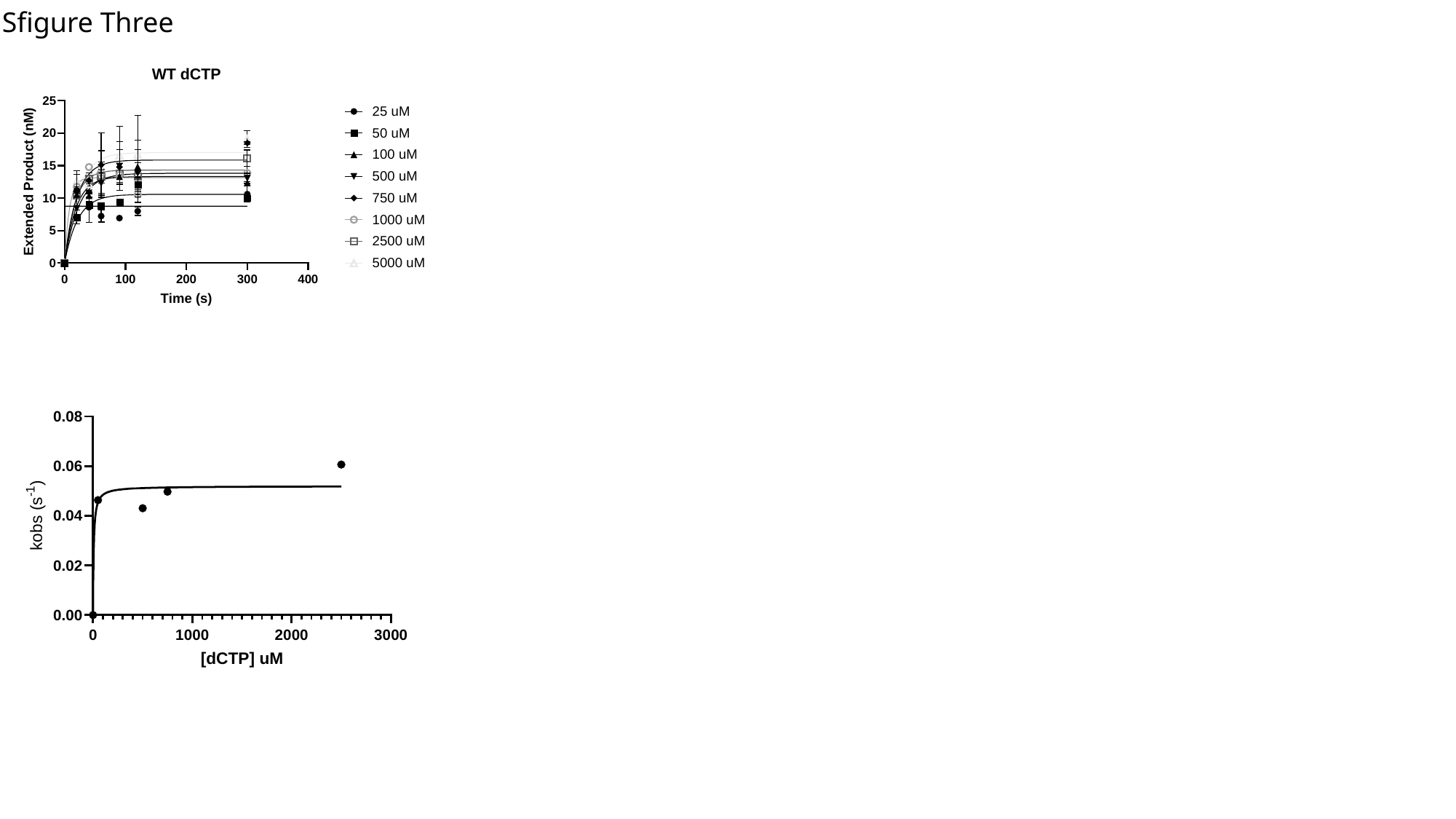

Sfigure Three

### Slide 4
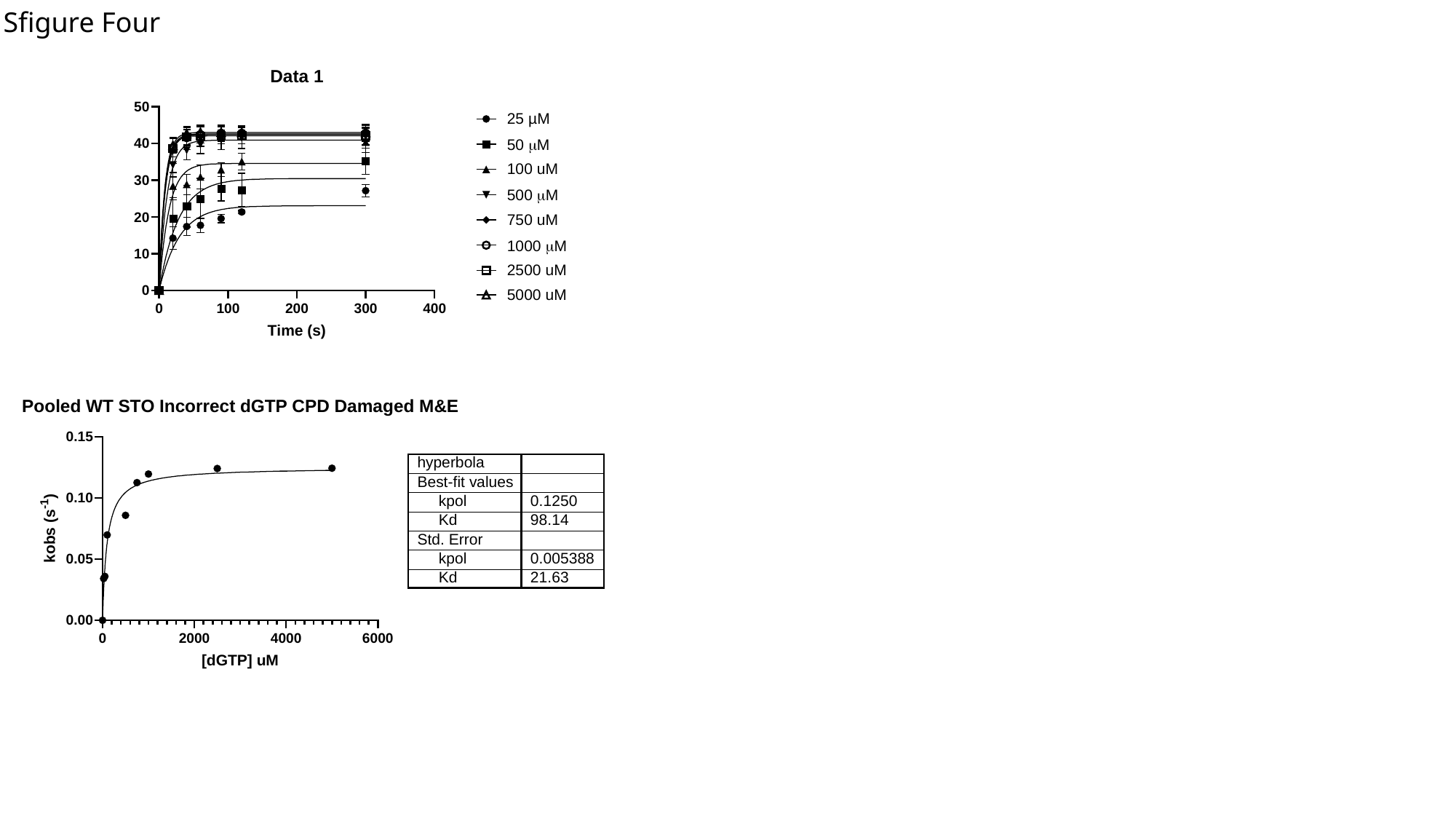

Sfigure Four

### Slide 5
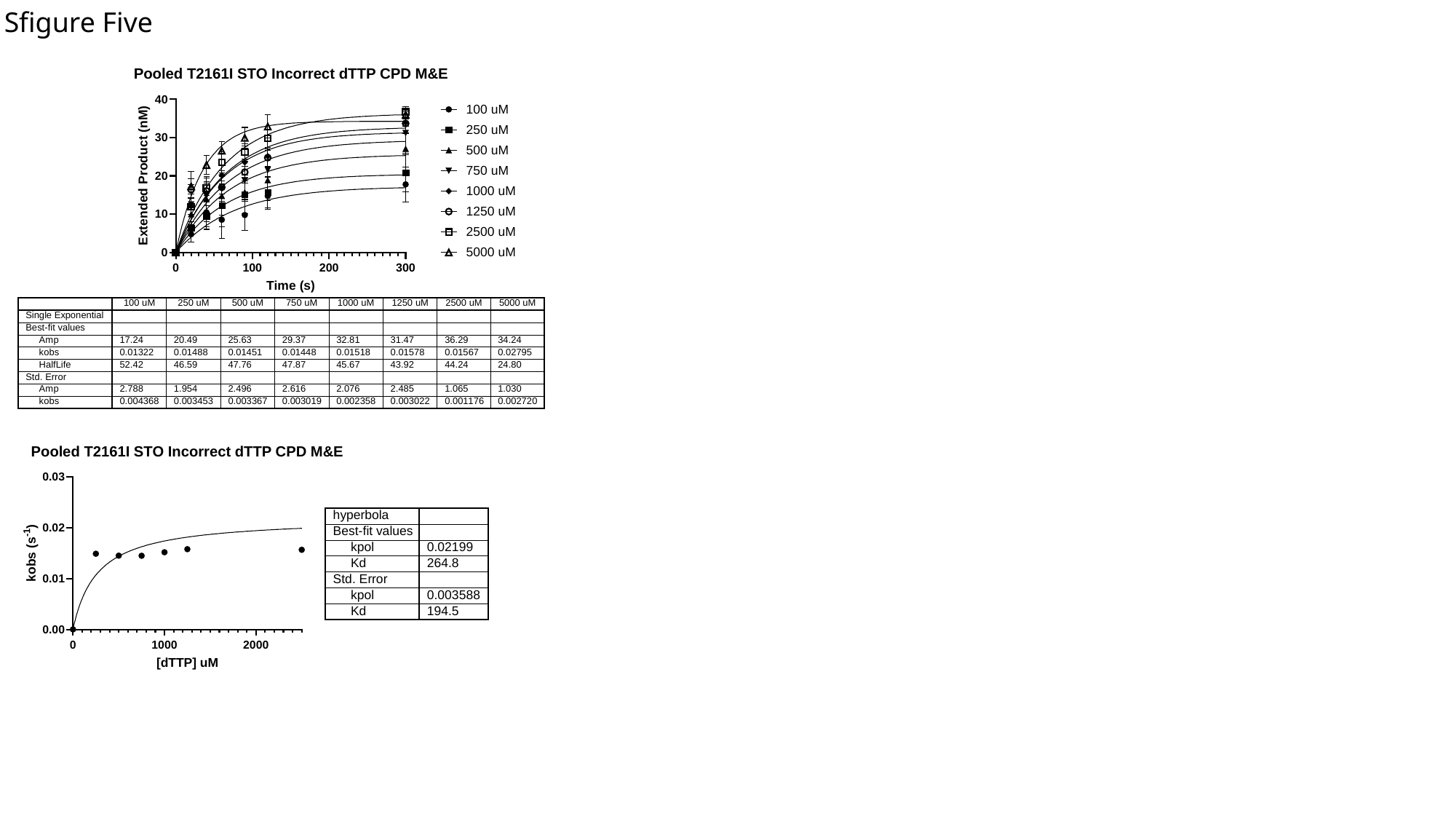

Sfigure Five

### Slide 6
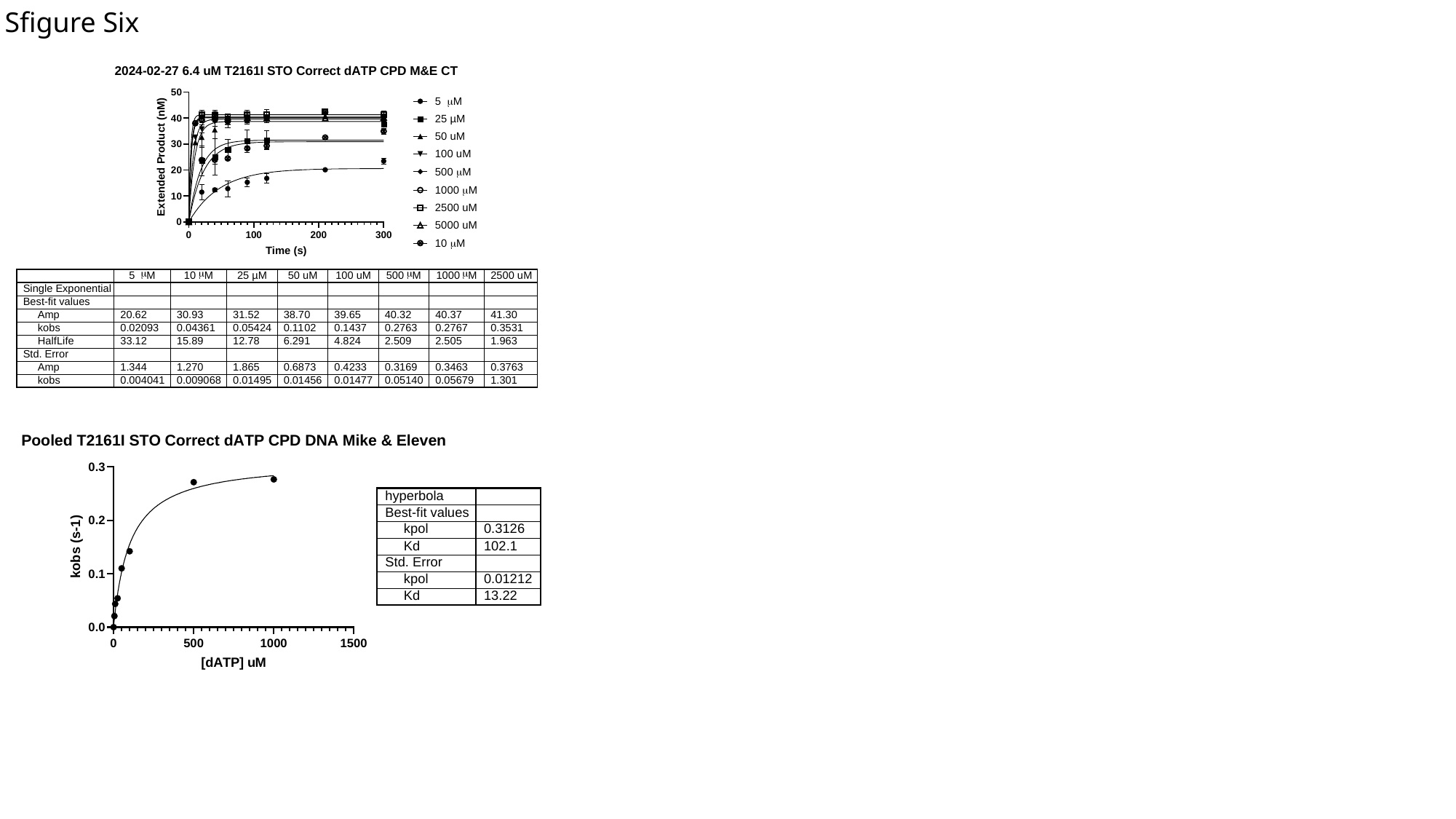

Sfigure Six

### Slide 7
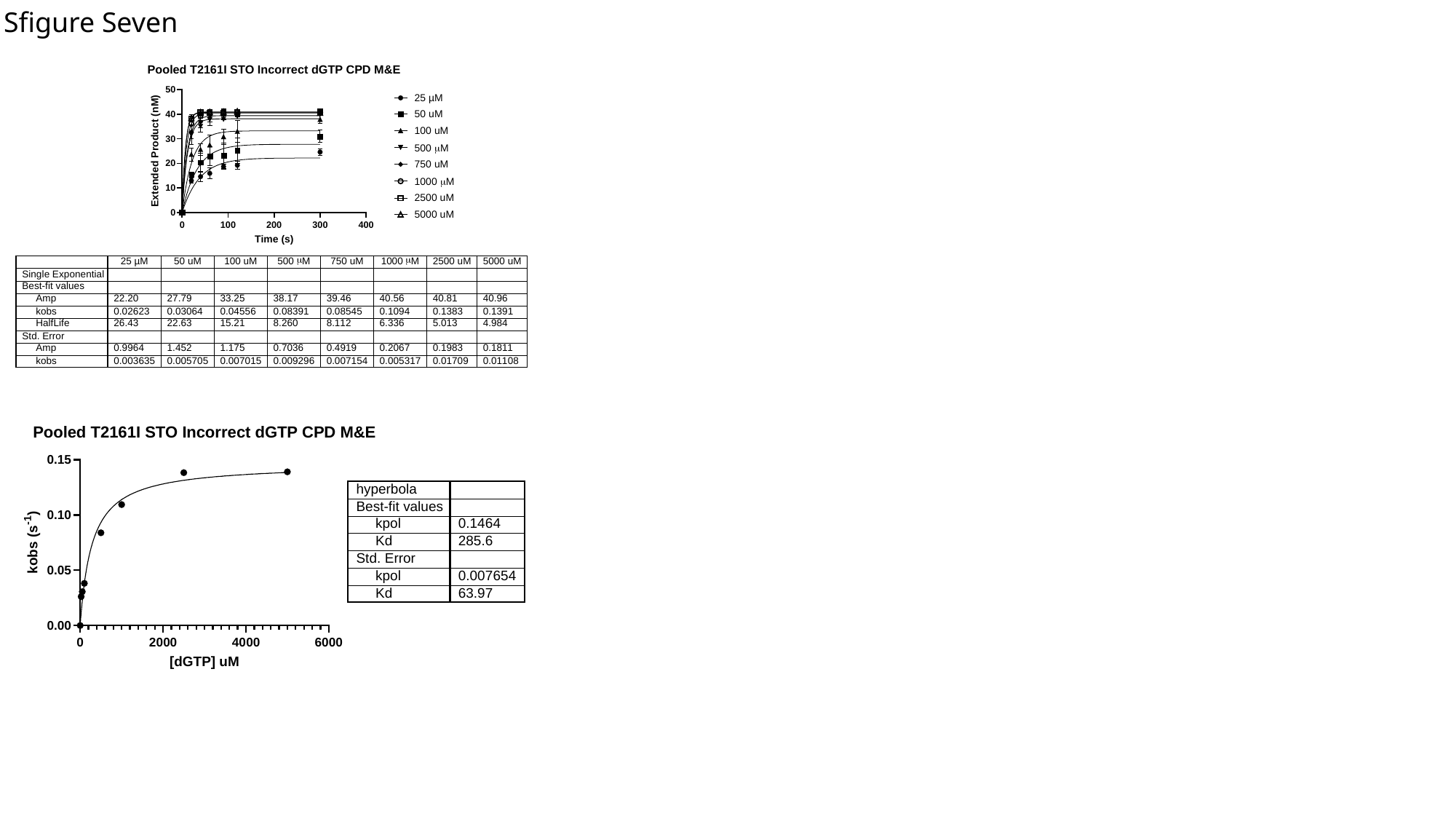

Sfigure Seven

### Slide 8
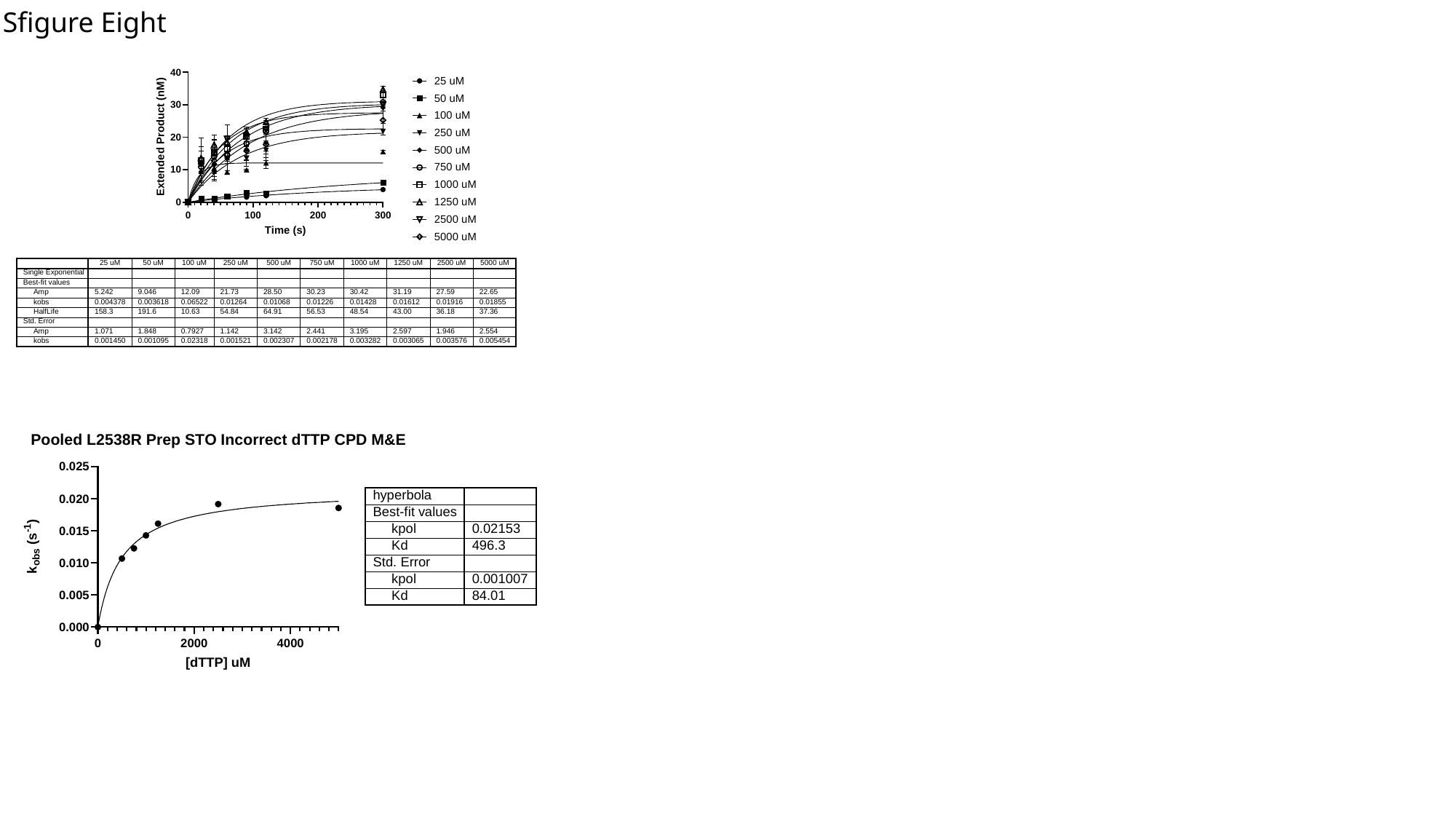

Sfigure Eight

### Slide 9
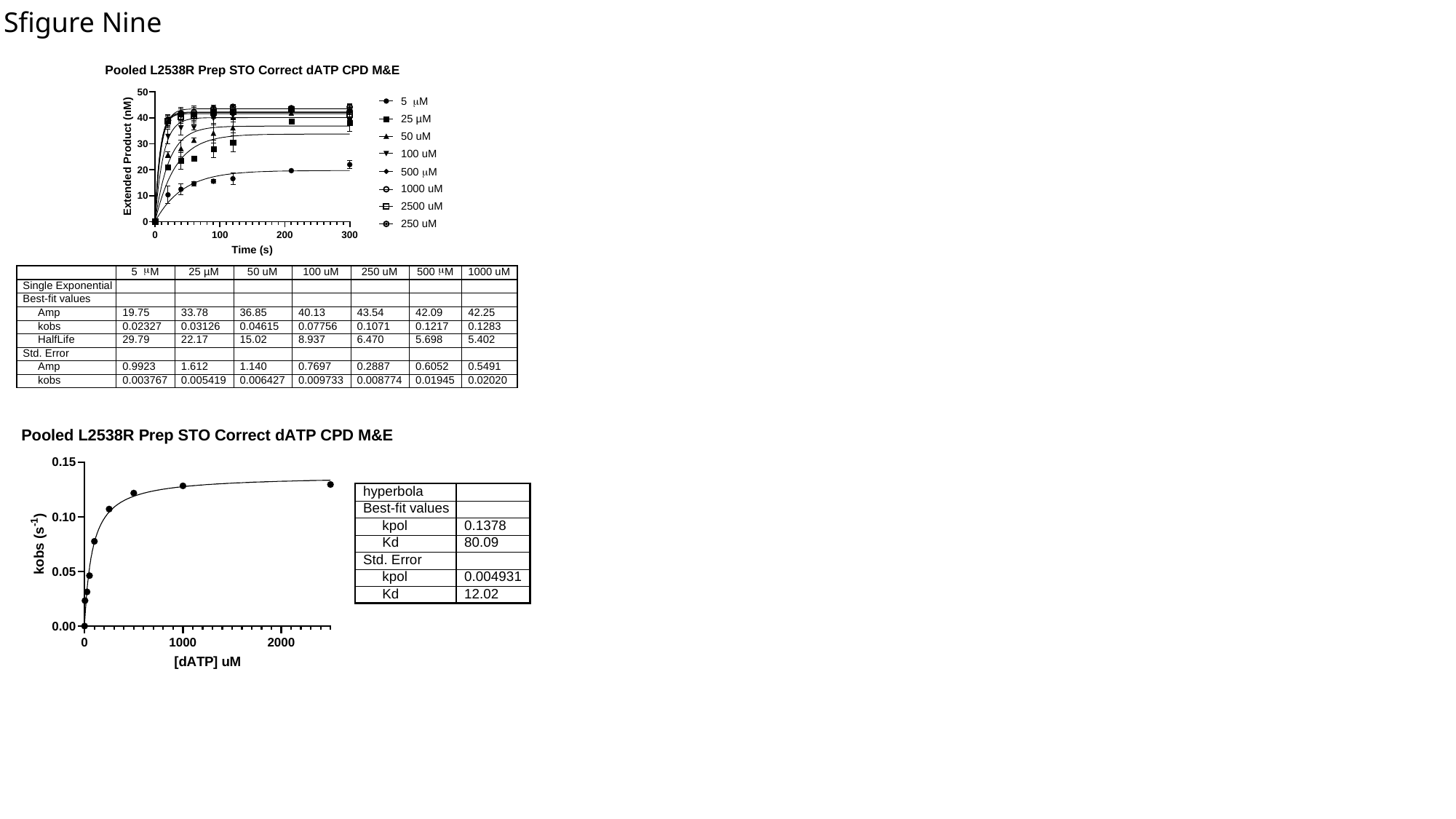

Sfigure Nine

### Slide 10
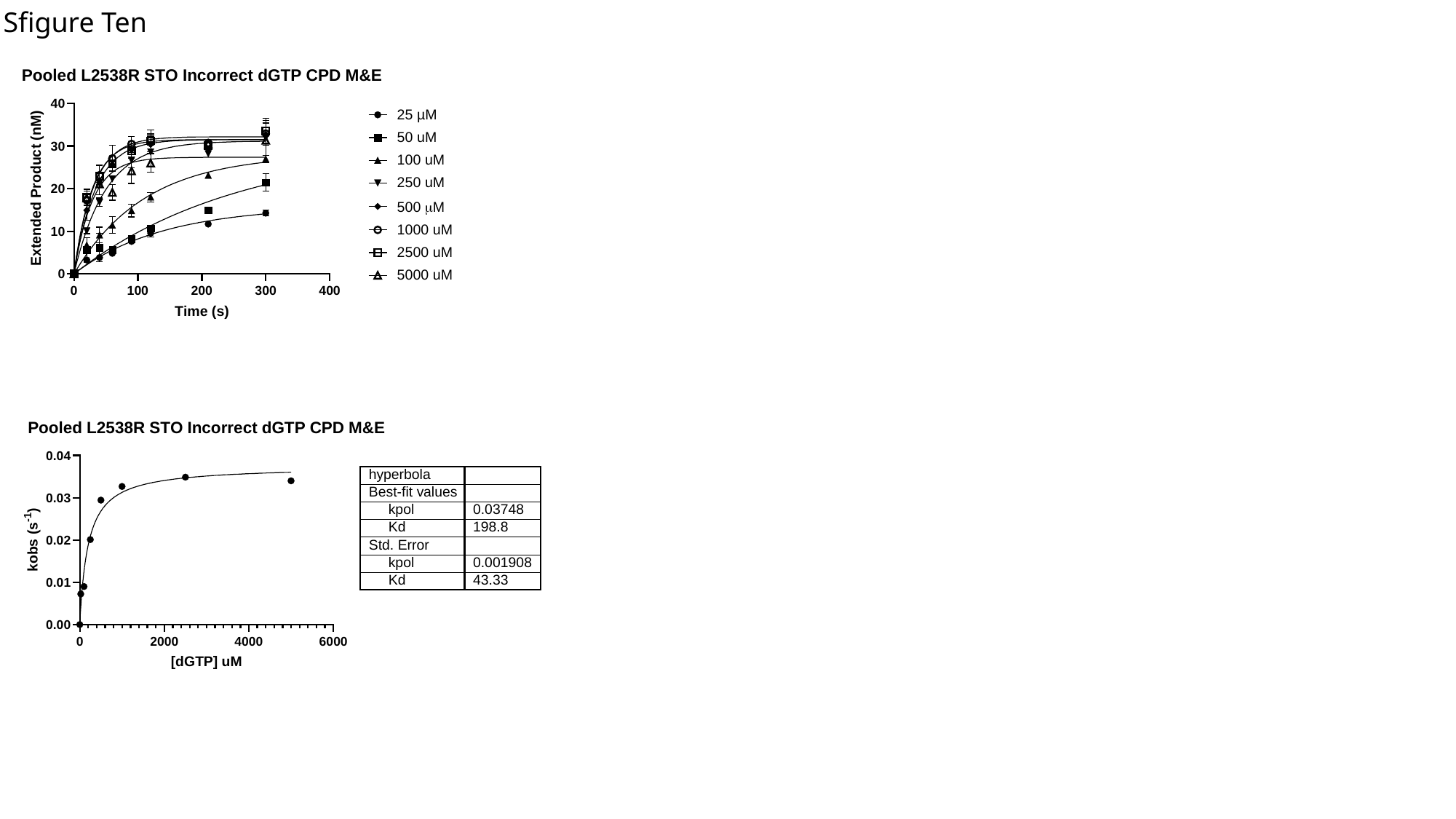

Sfigure Ten

### Slide 11
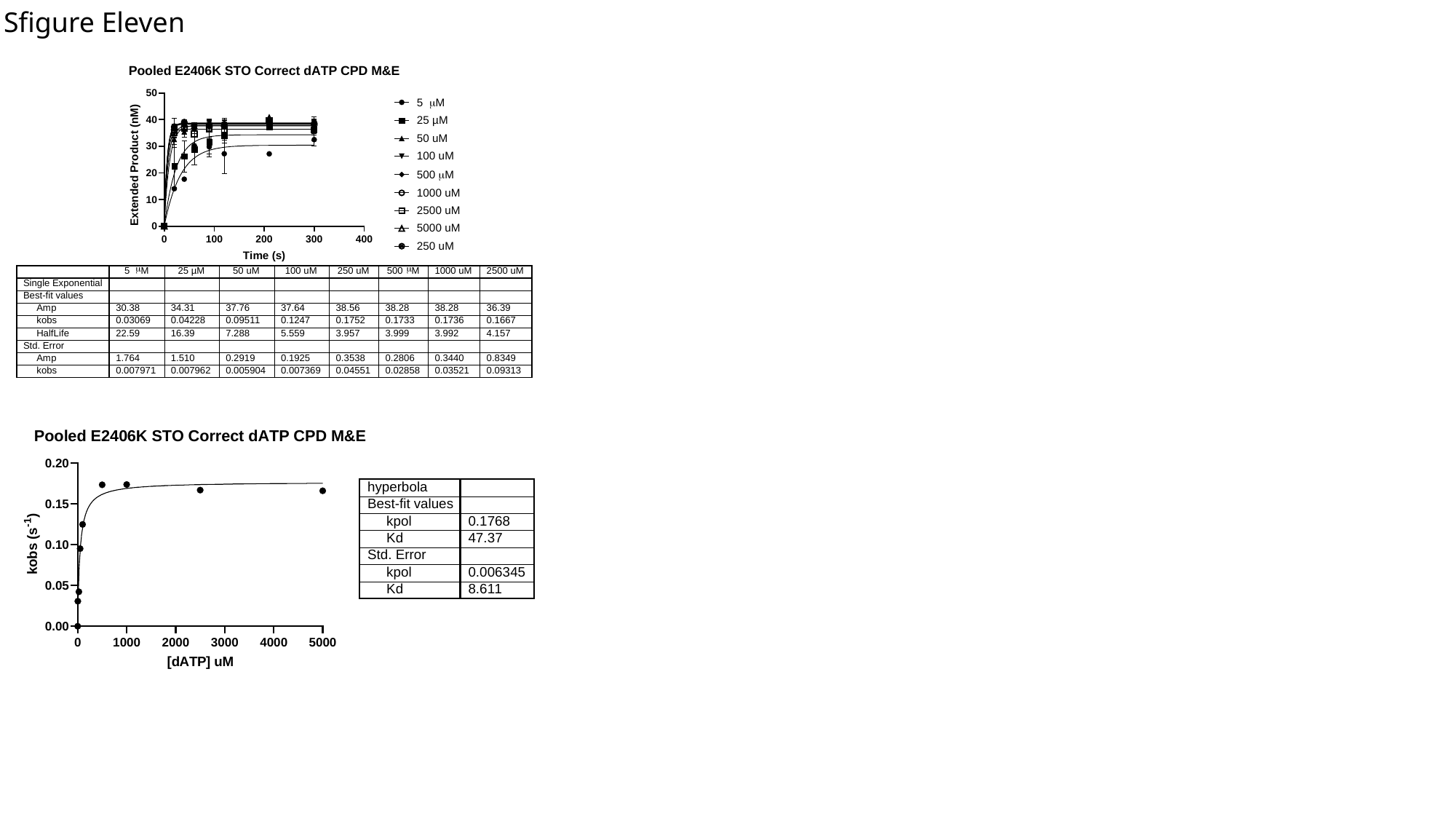

Sfigure Eleven

### Slide 12
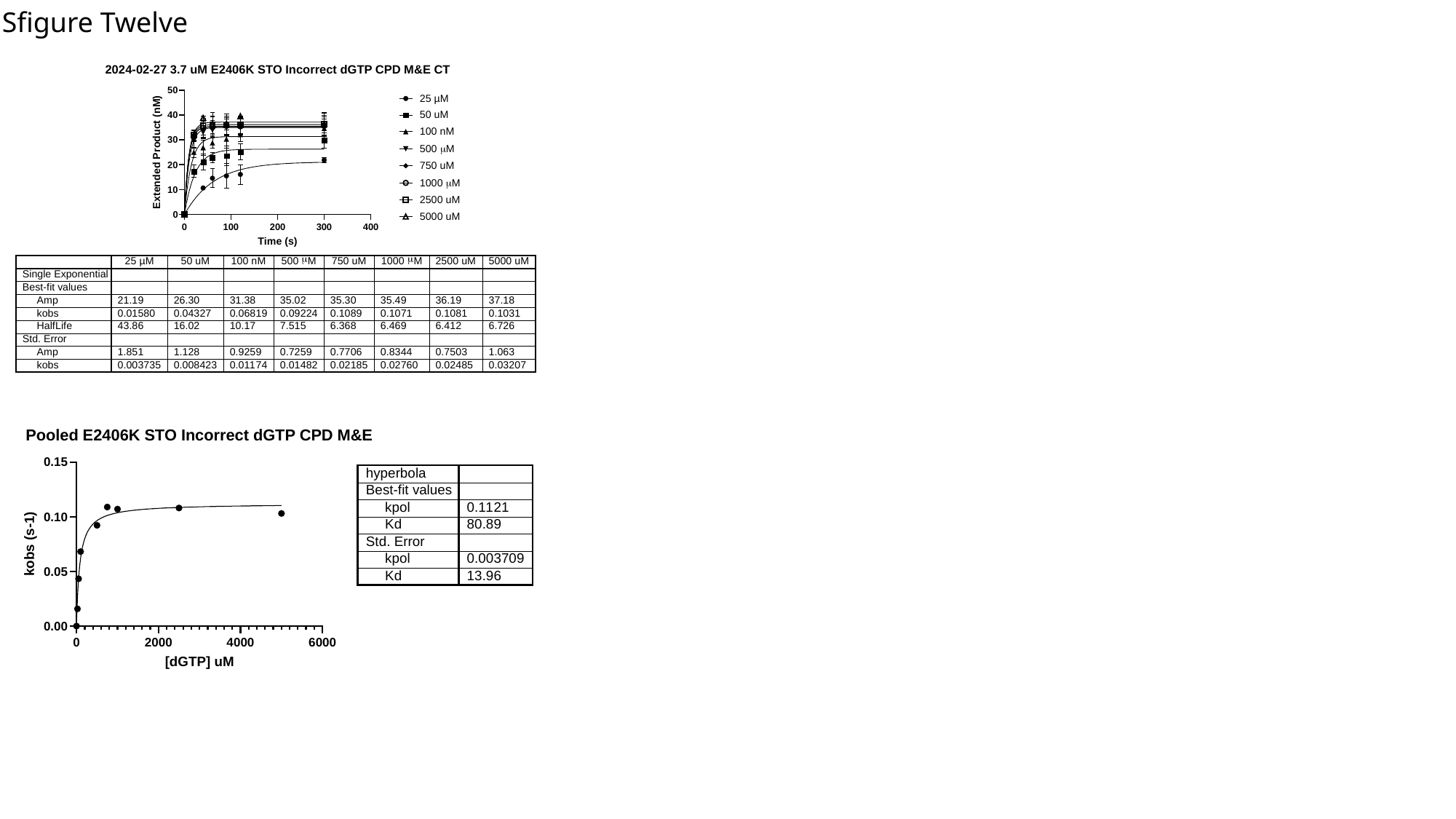

Sfigure Twelve

### Slide 13
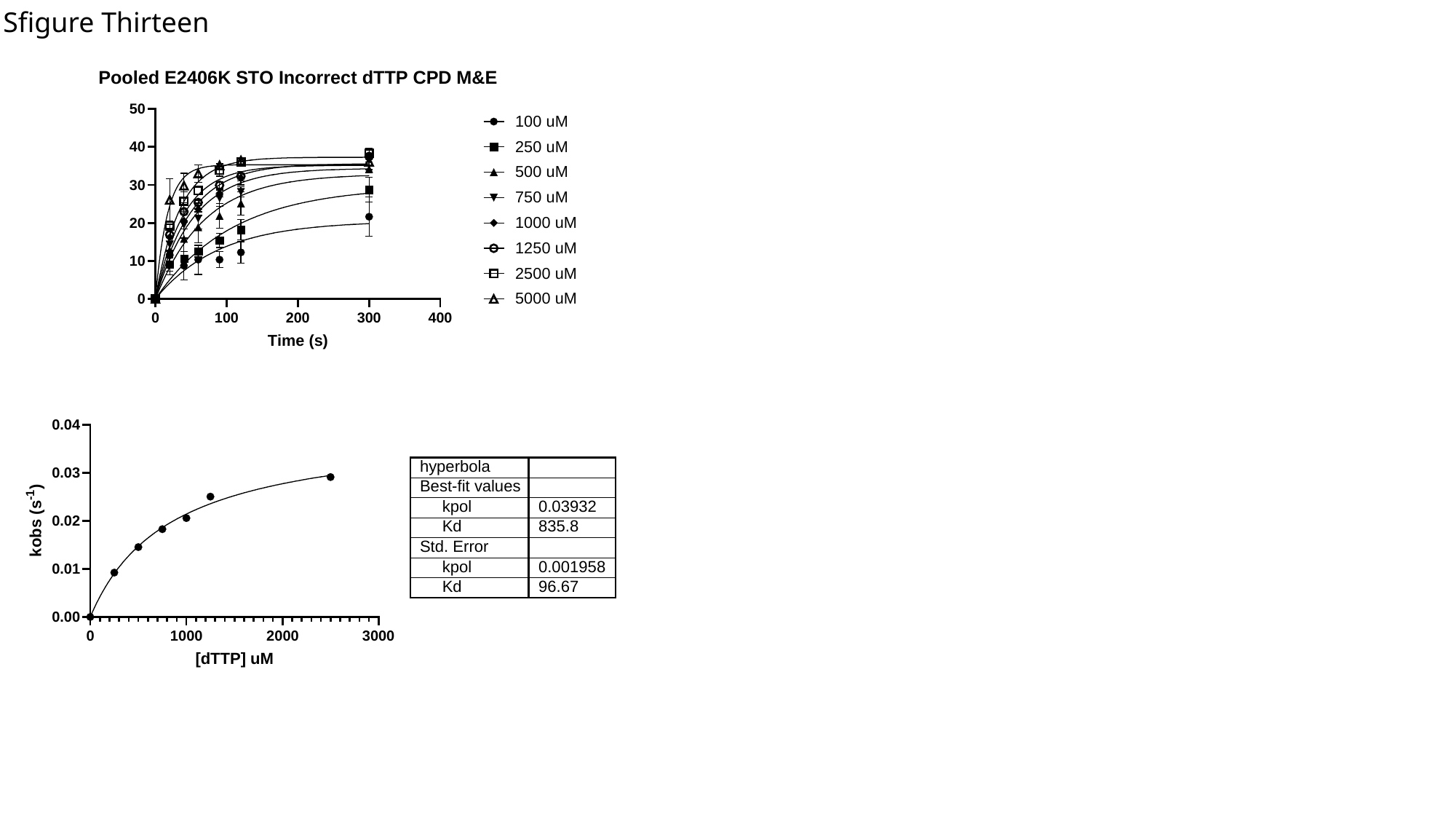

Sfigure Thirteen

### Slide 14
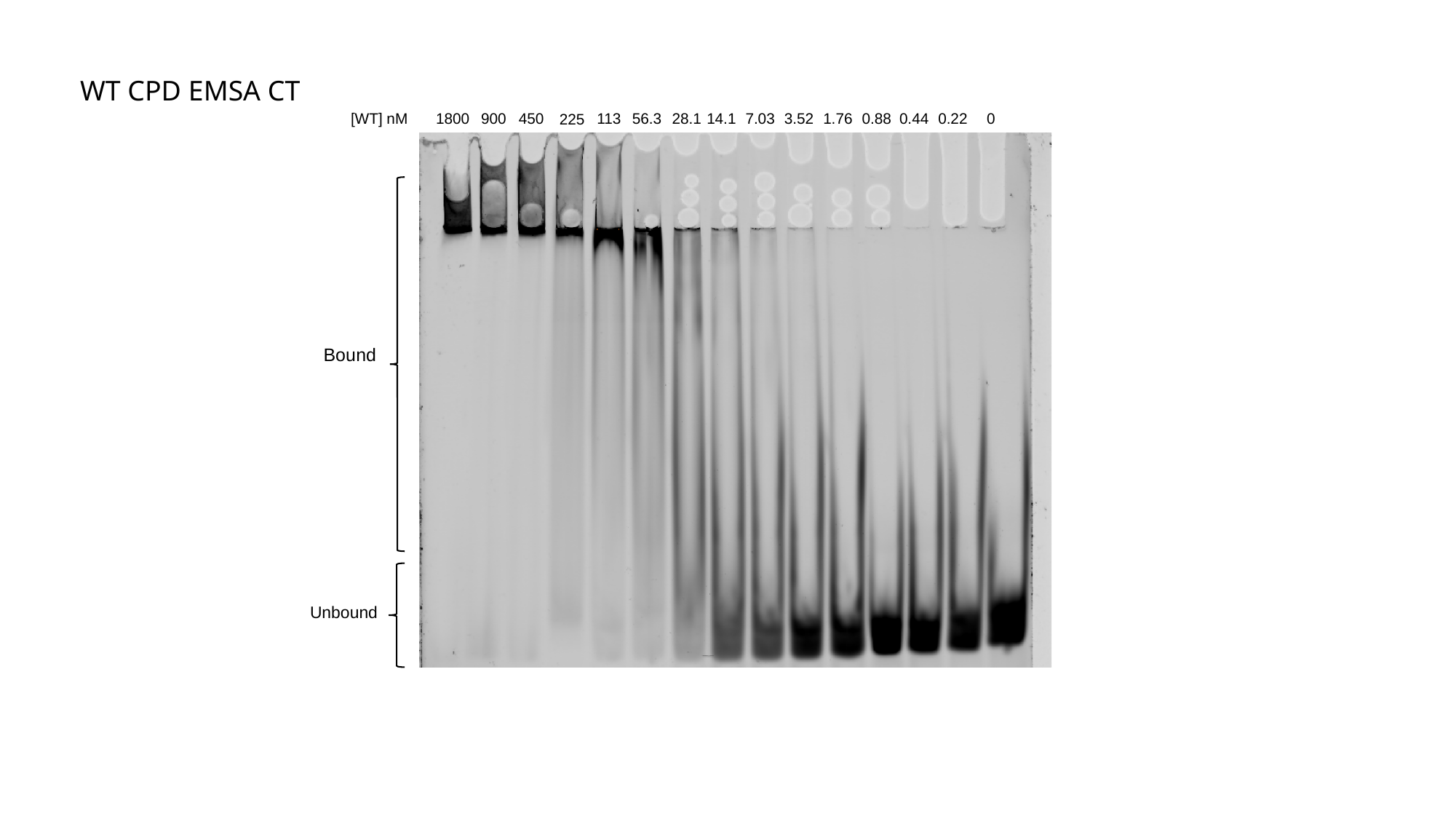

WT CPD EMSA CT
0
14.1
7.03
3.52
1.76
0.88
0.44
0.22
56.3
28.1
[WT] nM
1800
900
450
113
225
Bound
Unbound

### Slide 15
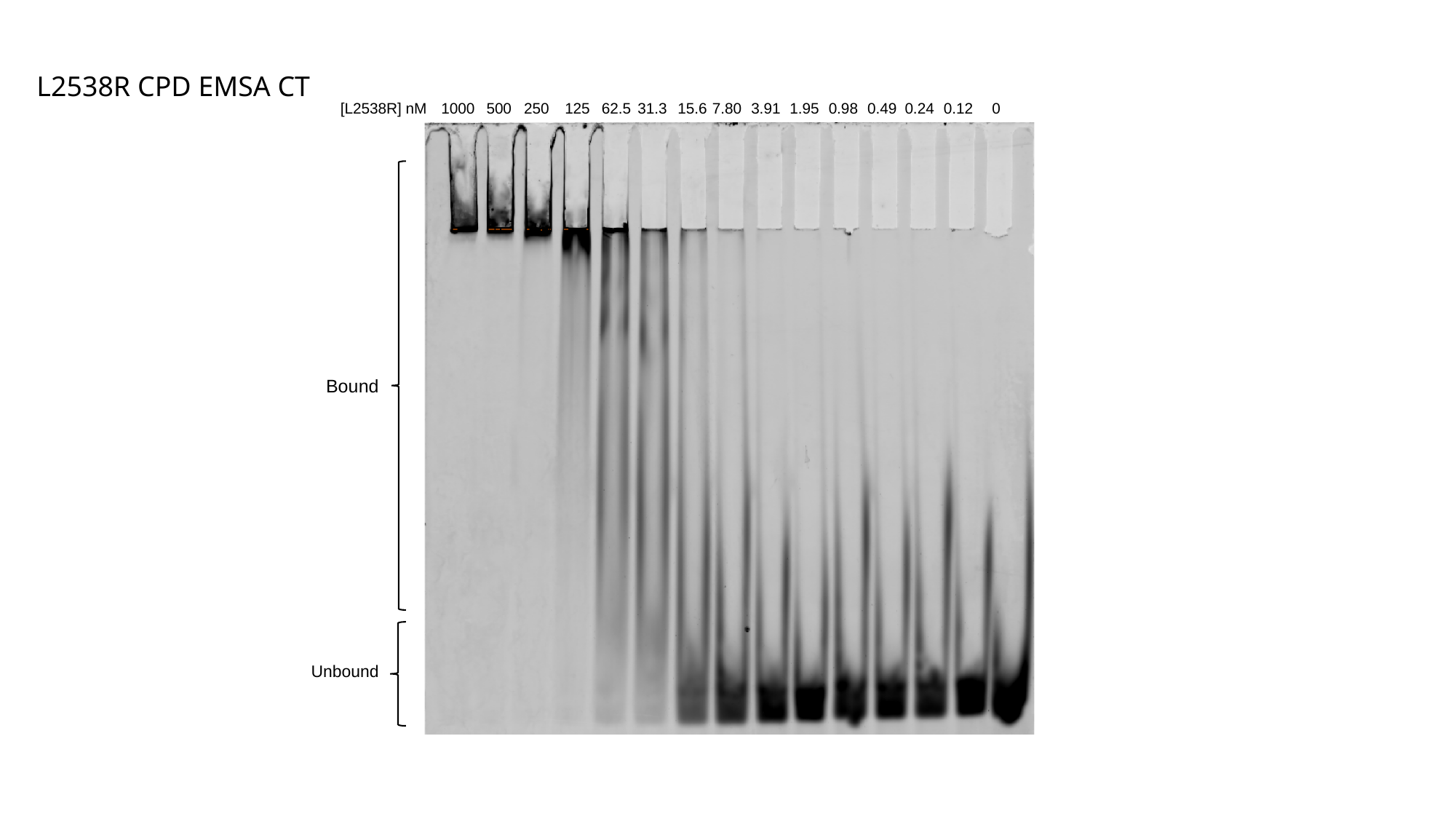

L2538R CPD EMSA CT
[L2538R] nM
0
7.80
3.91
1.95
0.98
0.49
0.24
0.12
31.3
15.6
1000
500
250
62.5
125
Bound
Unbound

### Slide 16
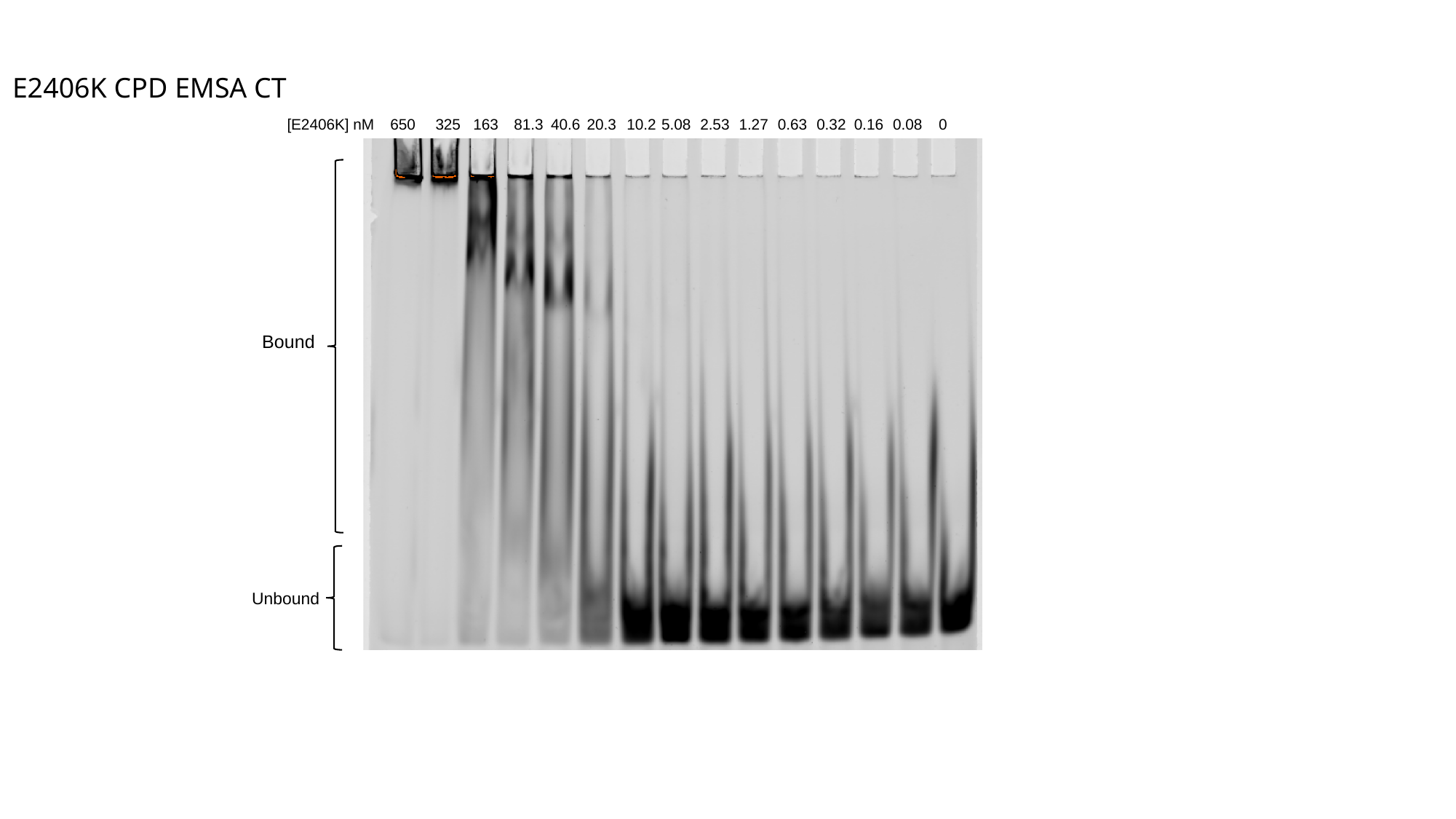

E2406K CPD EMSA CT
[E2406K] nM
5.08
2.53
1.27
0.63
0.32
0.16
0.08
0
20.3
10.2
650
325
163
40.6
81.3
Bound
Unbound

### Slide 17
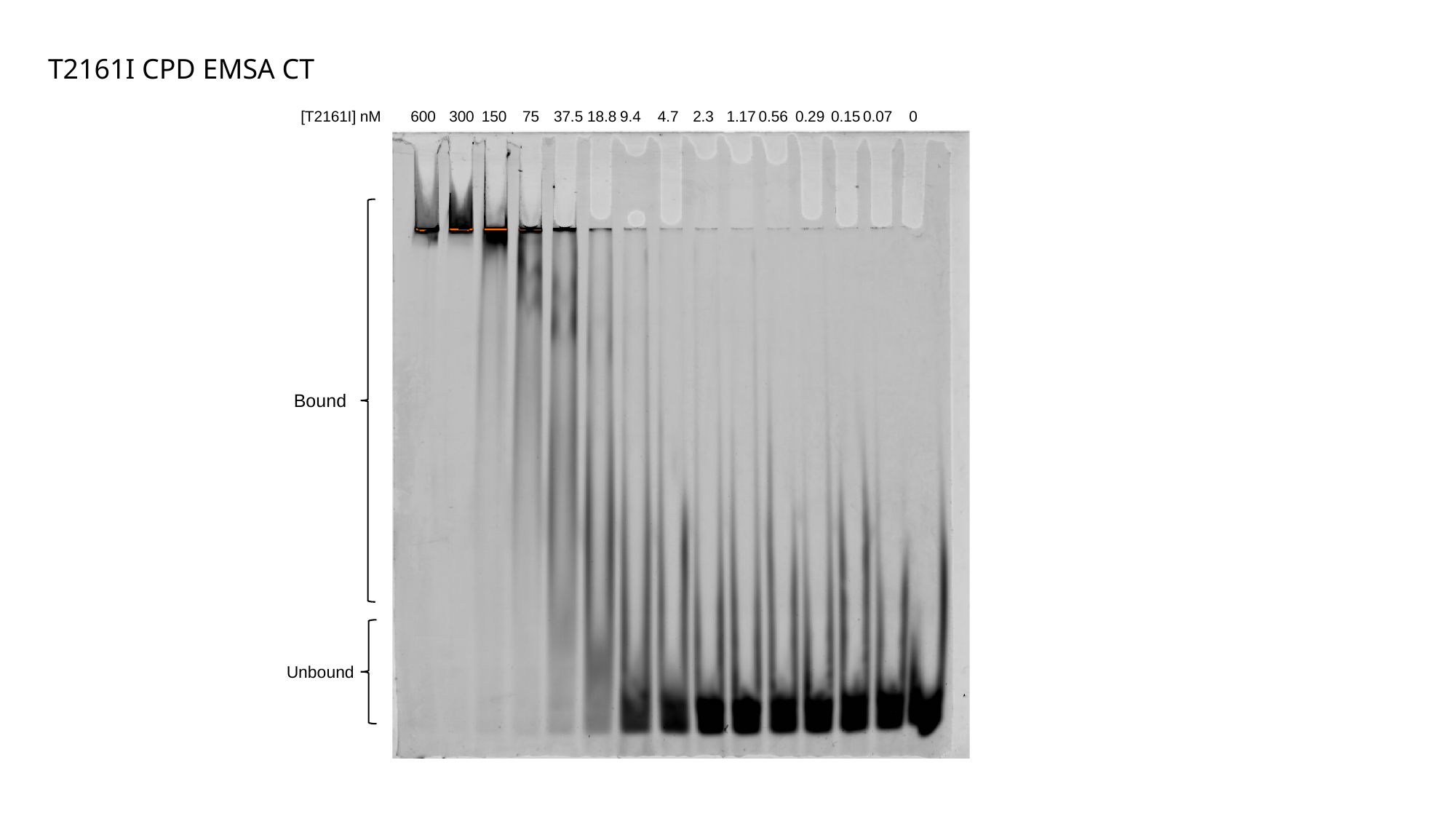

T2161I CPD EMSA CT
0.15
0.07
0
0.29
0.56
1.17
4.7
2.3
150
37.5
18.8
9.4
[T2161I] nM
75
600
300
Bound
Unbound

### Slide 18
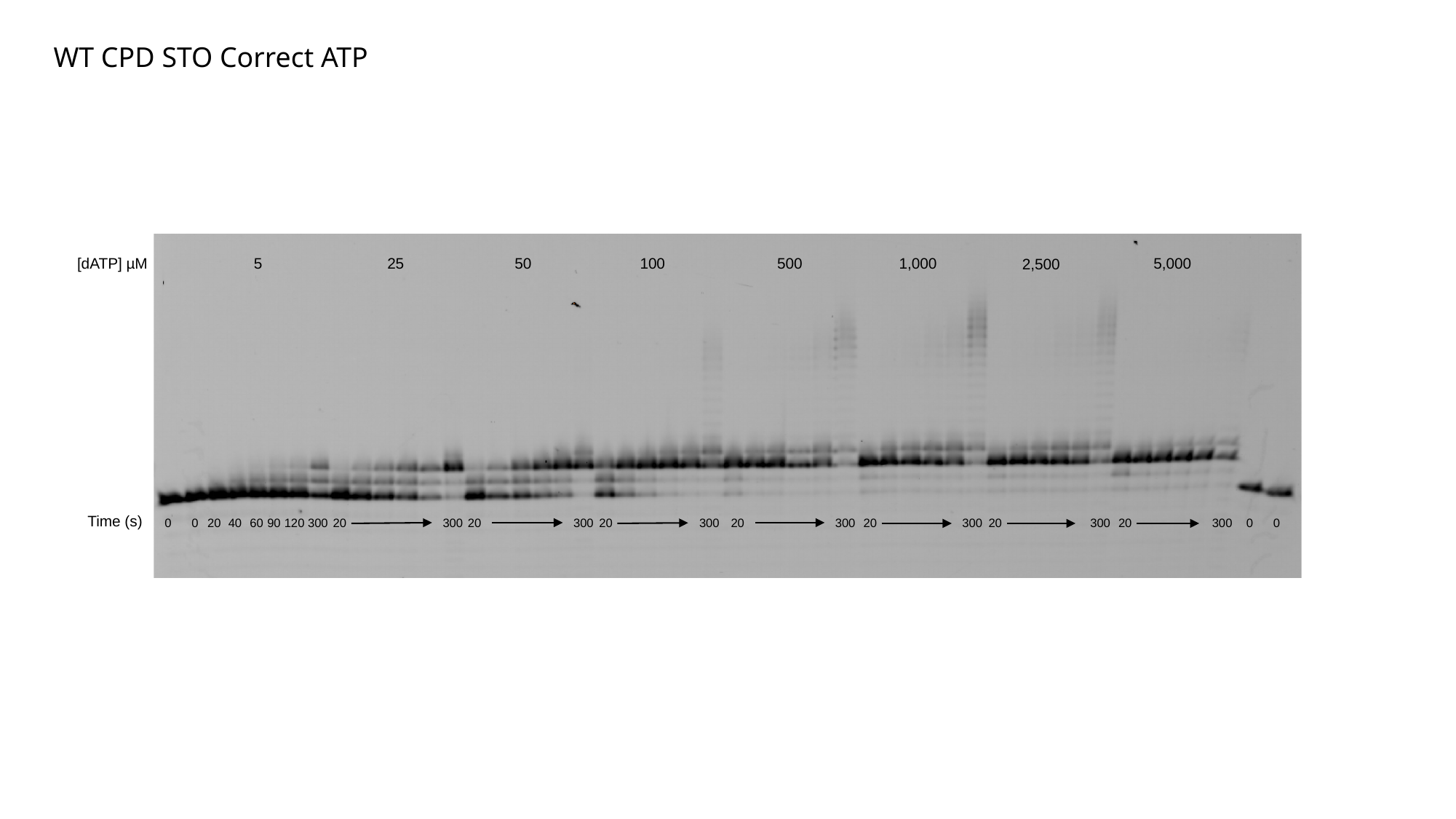

WT CPD STO Correct ATP
5,000
500
1,000
100
[dATP] µM
5
25
50
2,500
Time (s)
0
0
20
40
60
90
120
300
20
300
20
300
20
300
20
300
20
300
20
300
20
300
0
0

### Slide 19
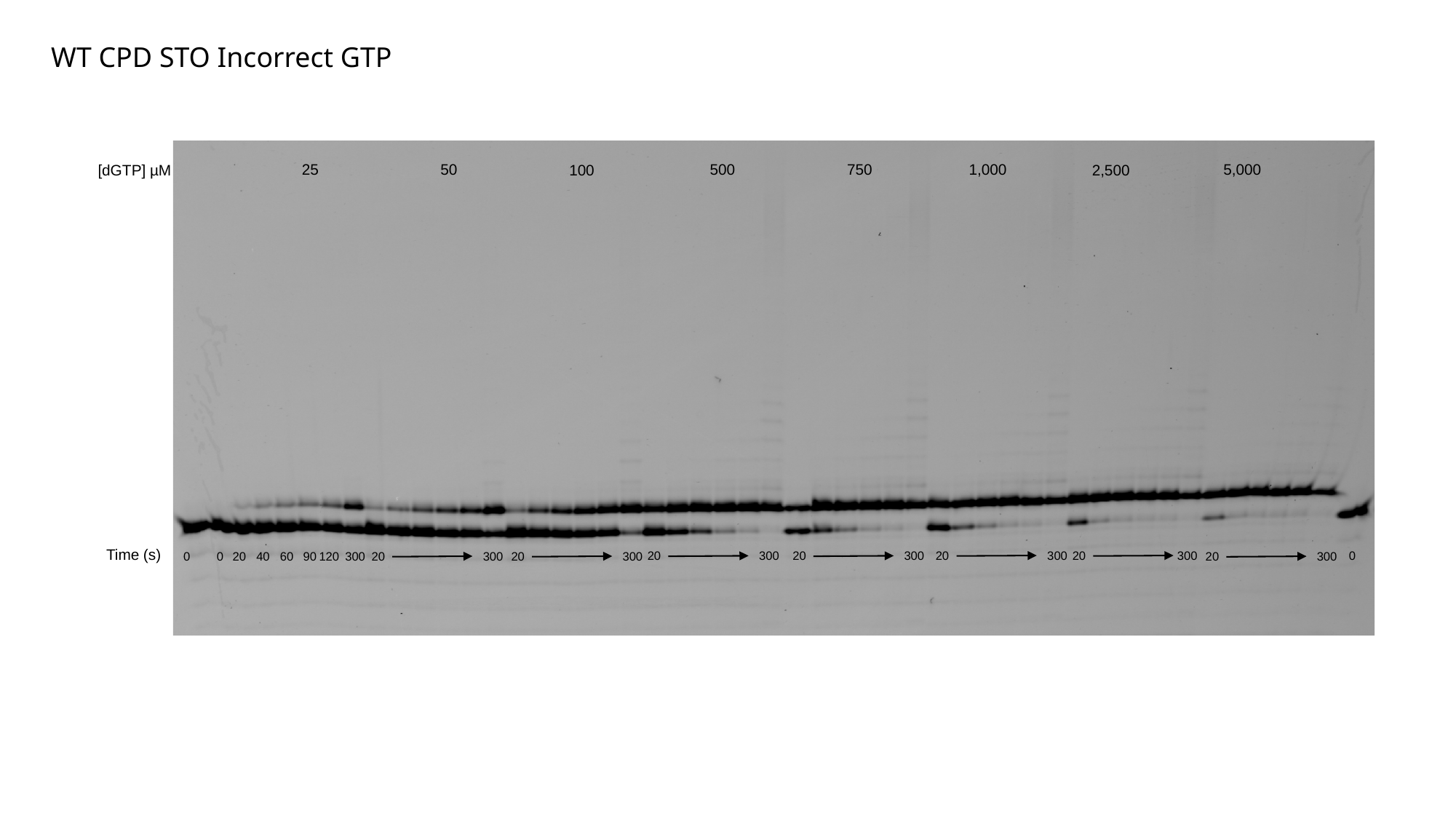

WT CPD STO Incorrect GTP
25
50
5,000
750
1,000
500
[dGTP] µM
100
2,500
Time (s)
20
300
20
300
0
20
300
20
300
0
0
20
40
60
90
120
300
20
300
20
300
20
300

### Slide 20
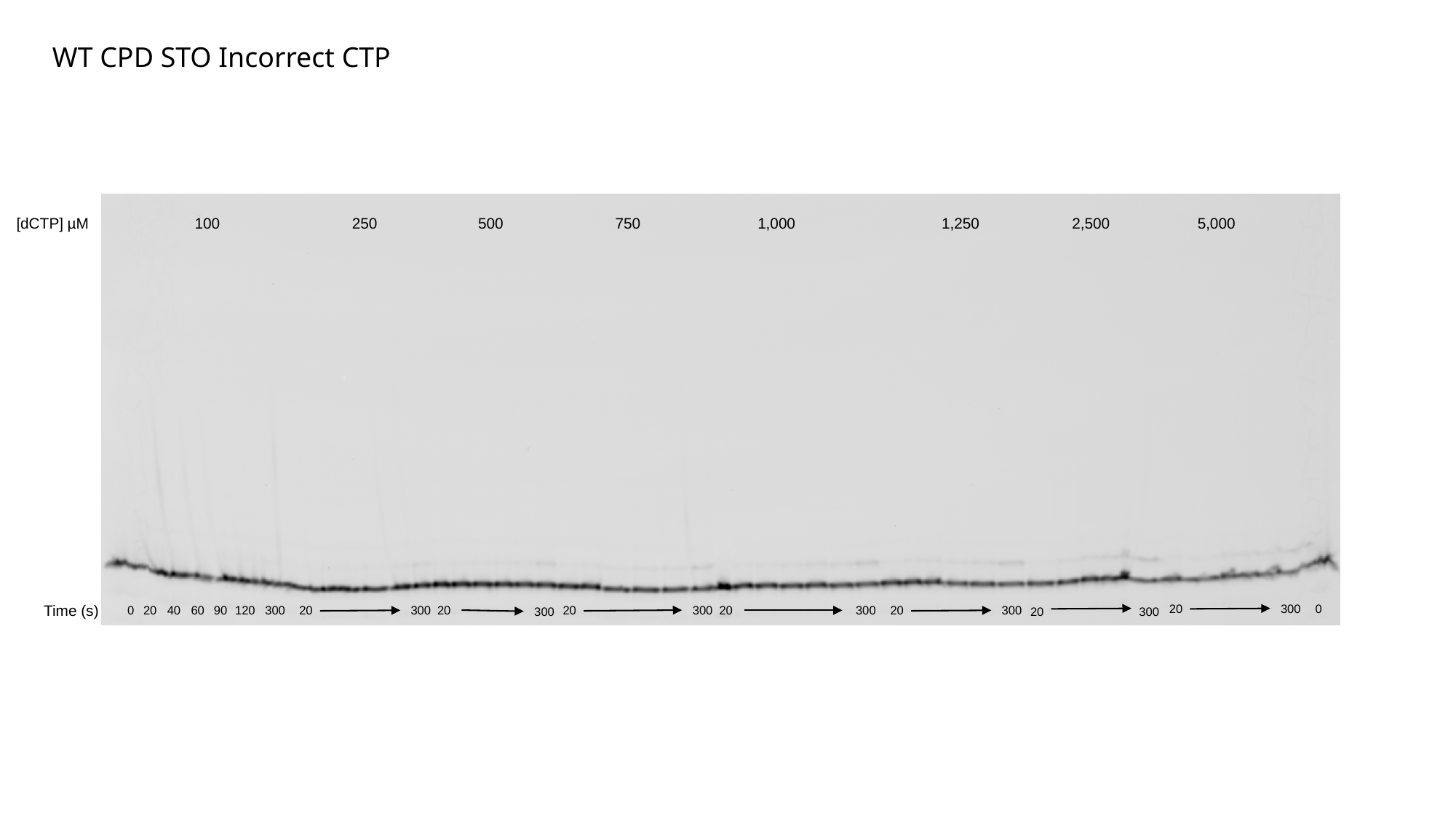

WT CPD STO Incorrect CTP
100
250
500
750
1,000
1,250
2,500
[dCTP] µM
5,000
20
300
0
Time (s)
0
20
40
60
90
120
300
20
300
20
20
300
20
300
20
300
300
20
300

### Slide 21
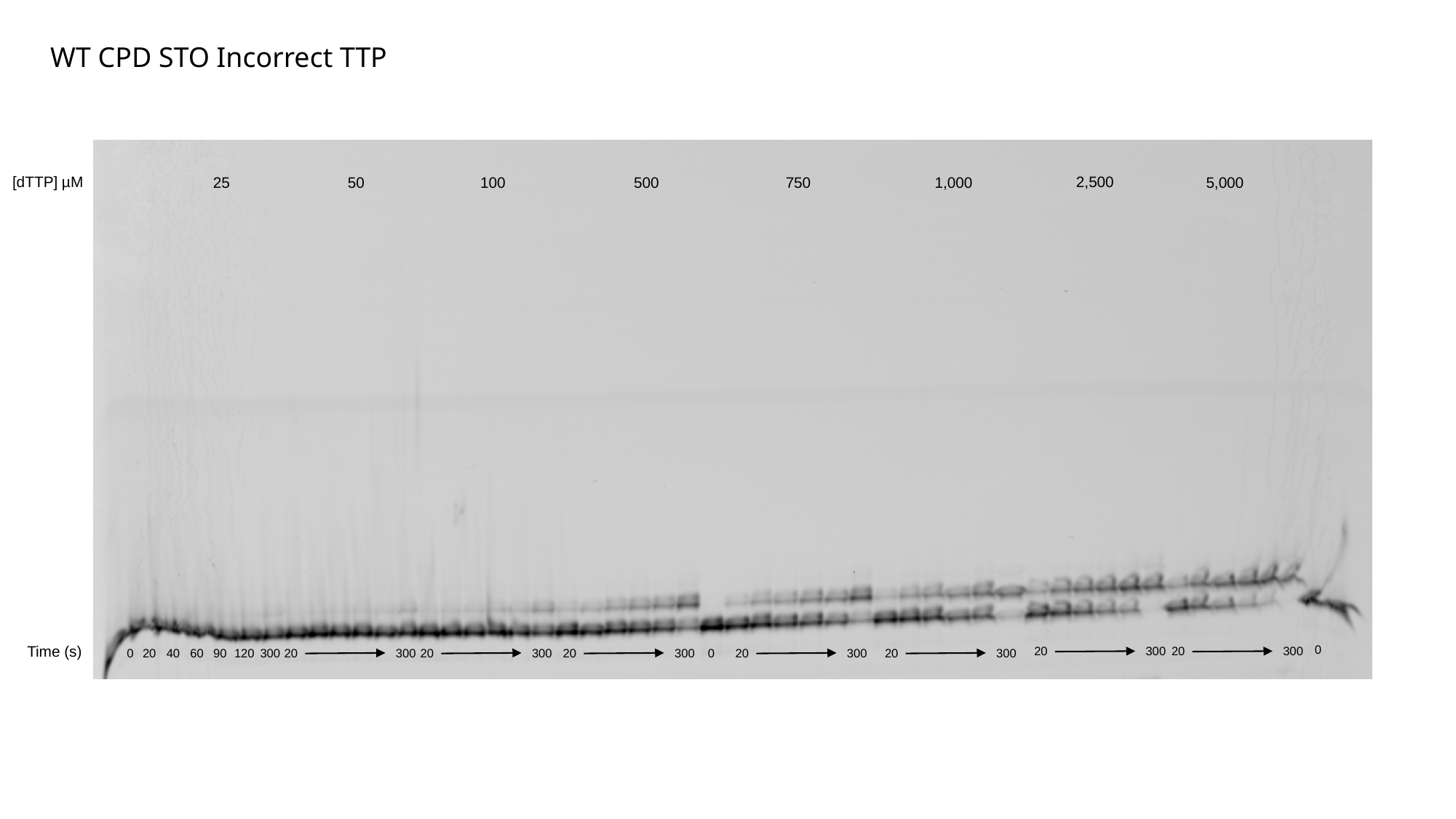

WT CPD STO Incorrect TTP
[dTTP] µM
2,500
25
50
100
500
750
1,000
5,000
Time (s)
0
20
300
20
300
0
20
40
60
90
120
300
20
300
20
300
20
300
0
20
300
20
300

### Slide 22
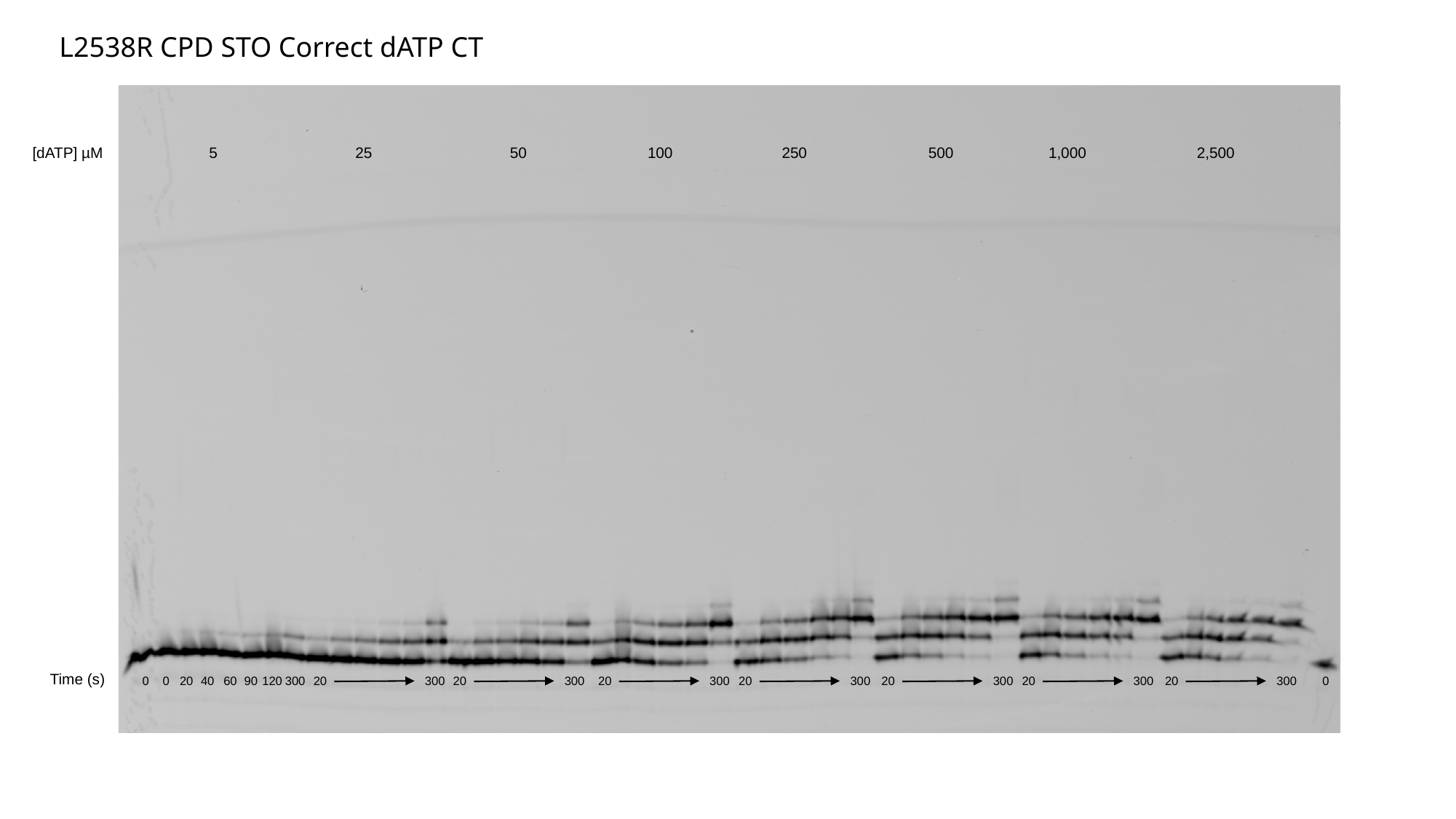

L2538R CPD STO Correct dATP CT
25
50
100
250
500
1,000
2,500
[dATP] µM
5
Time (s)
0
0
20
40
60
90
120
300
20
300
20
300
20
300
20
300
20
300
20
300
20
300
0

### Slide 23
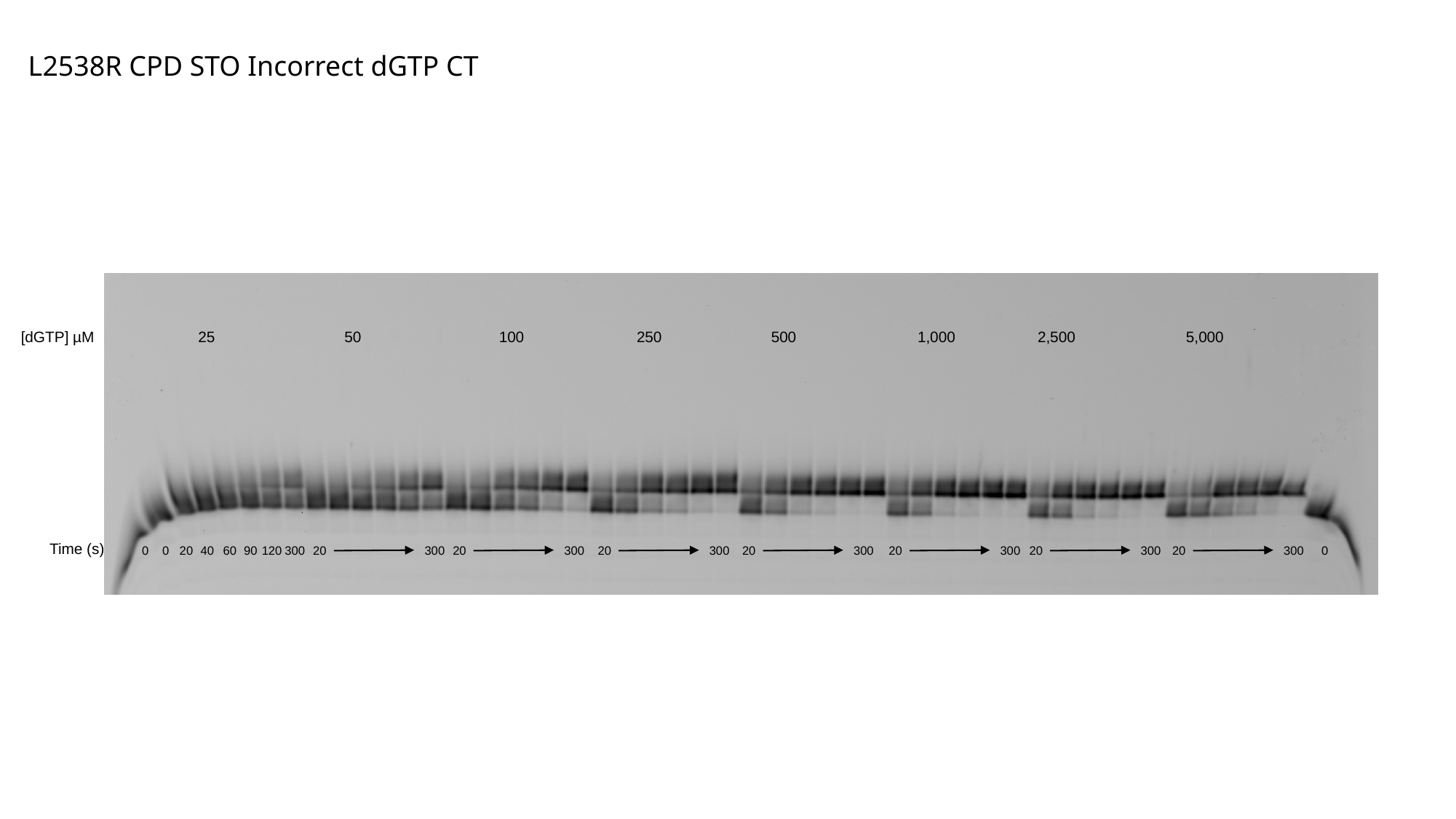

L2538R CPD STO Incorrect dGTP CT
50
100
250
500
1,000
2,500
5,000
[dGTP] µM
25
Time (s)
0
0
20
40
60
90
120
300
20
300
20
300
20
300
20
300
20
300
20
300
20
300
0

### Slide 24
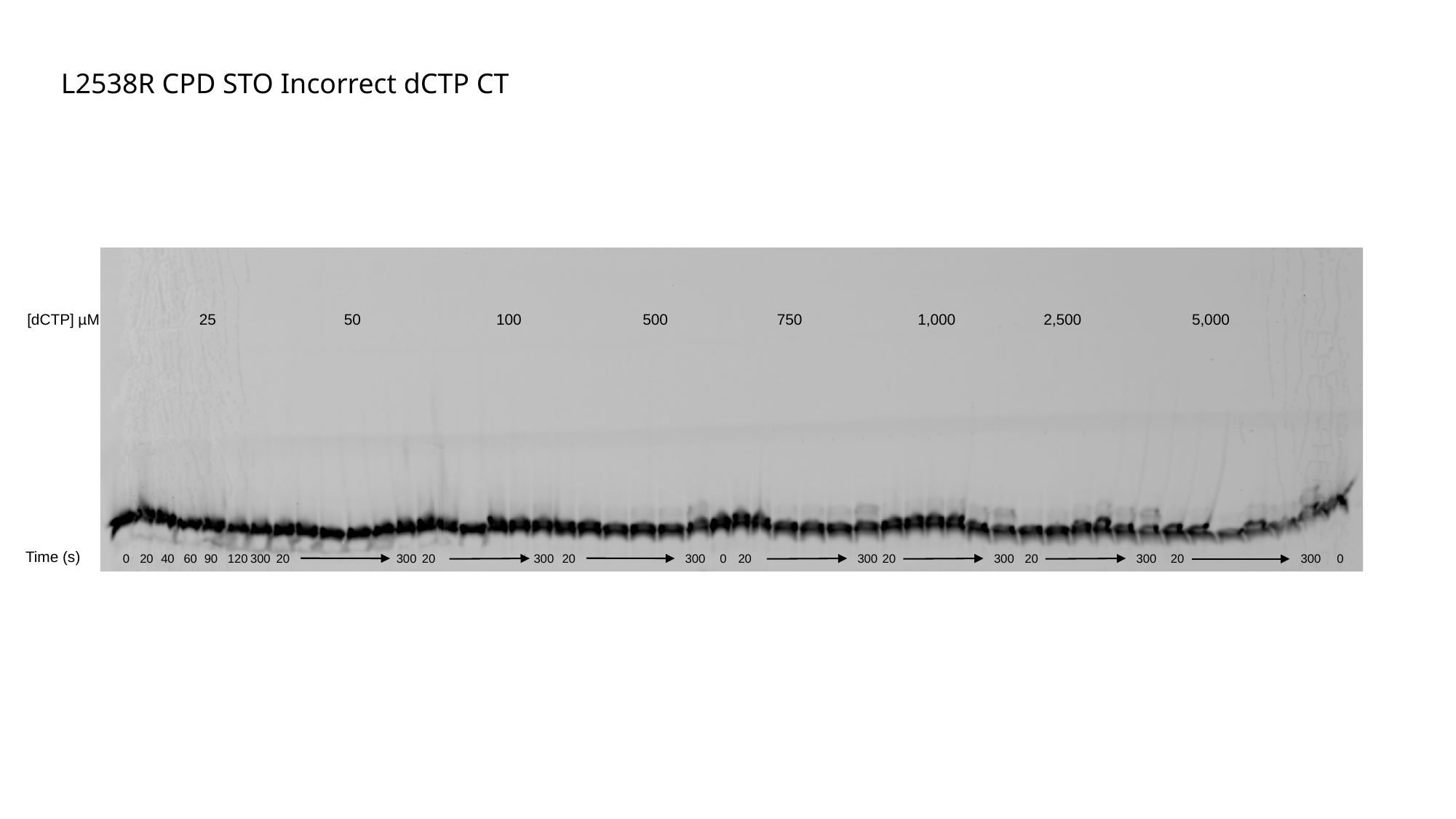

L2538R CPD STO Incorrect dCTP CT
50
100
500
1,000
750
2,500
5,000
[dCTP] µM
25
Time (s)
0
20
40
60
90
120
300
20
300
20
300
20
300
0
20
300
20
300
20
300
20
300
0

### Slide 25
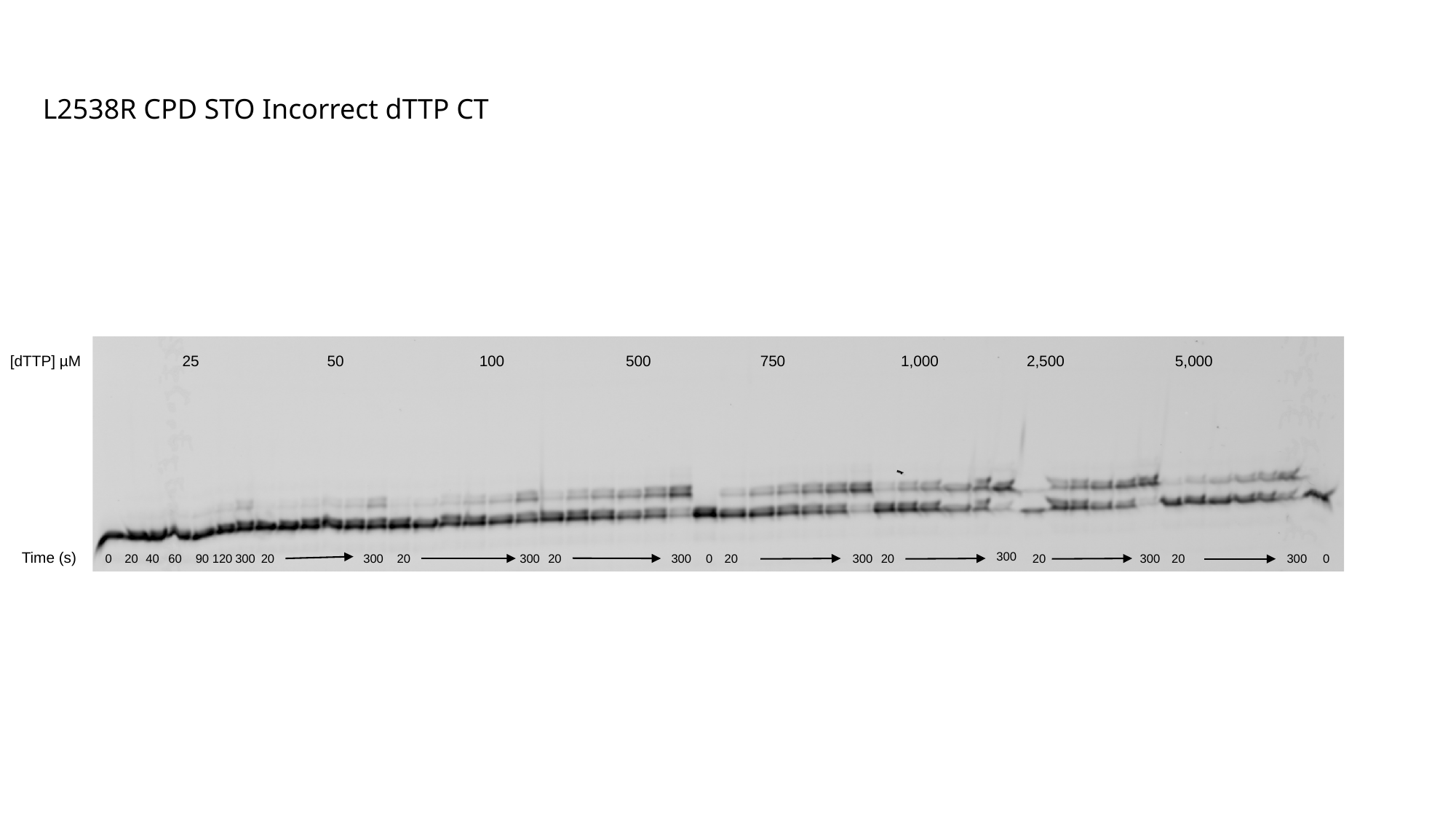

L2538R CPD STO Incorrect dTTP CT
50
100
500
1,000
750
2,500
5,000
[dTTP] µM
25
Time (s)
300
0
20
40
60
90
120
300
20
300
20
300
20
300
0
20
300
20
20
300
20
300
0

### Slide 26
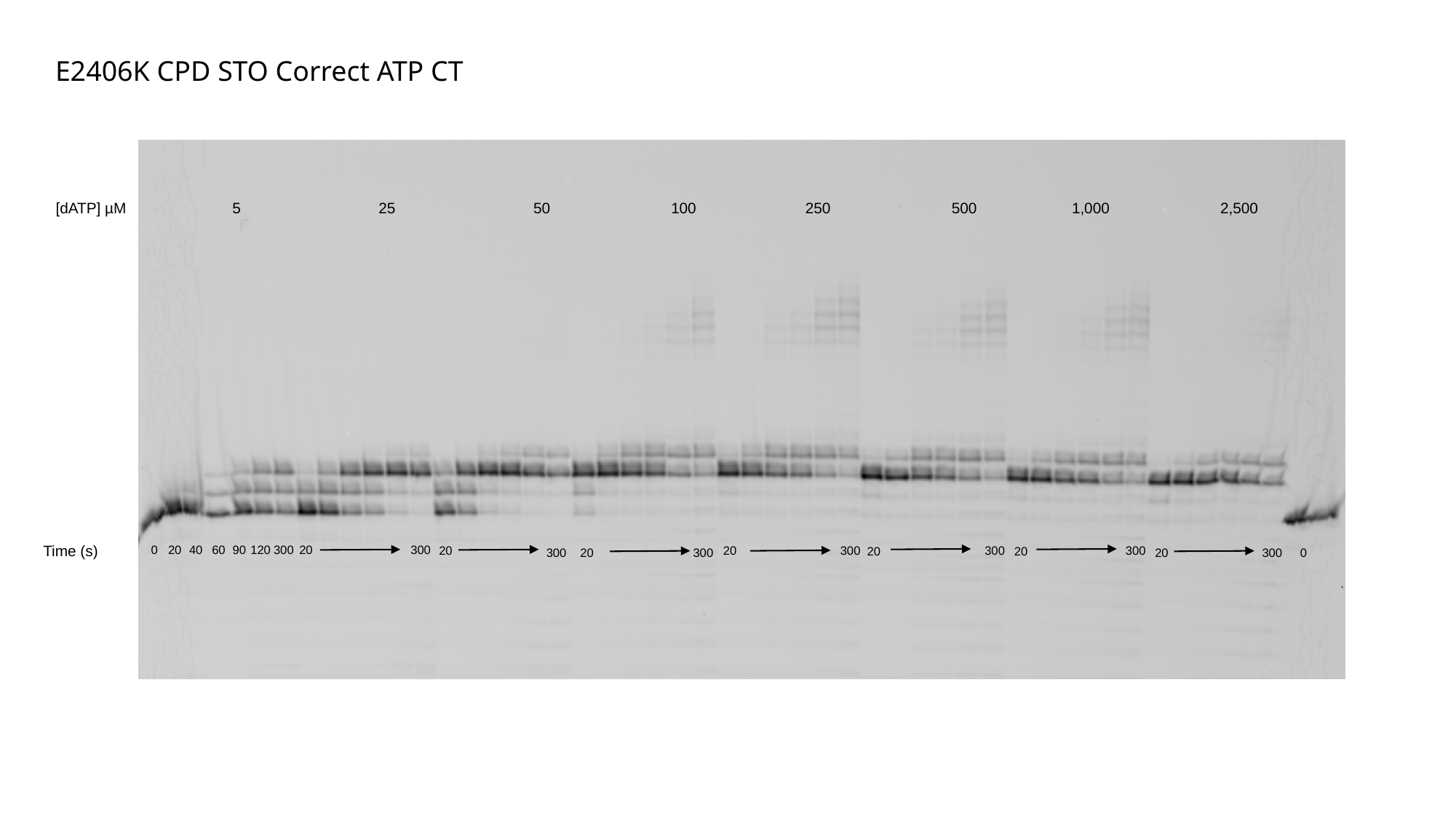

E2406K CPD STO Correct ATP CT
25
50
100
250
500
1,000
2,500
[dATP] µM
5
Time (s)
0
20
40
60
90
120
300
20
300
300
300
300
20
20
20
20
300
20
300
20
300
0

### Slide 27
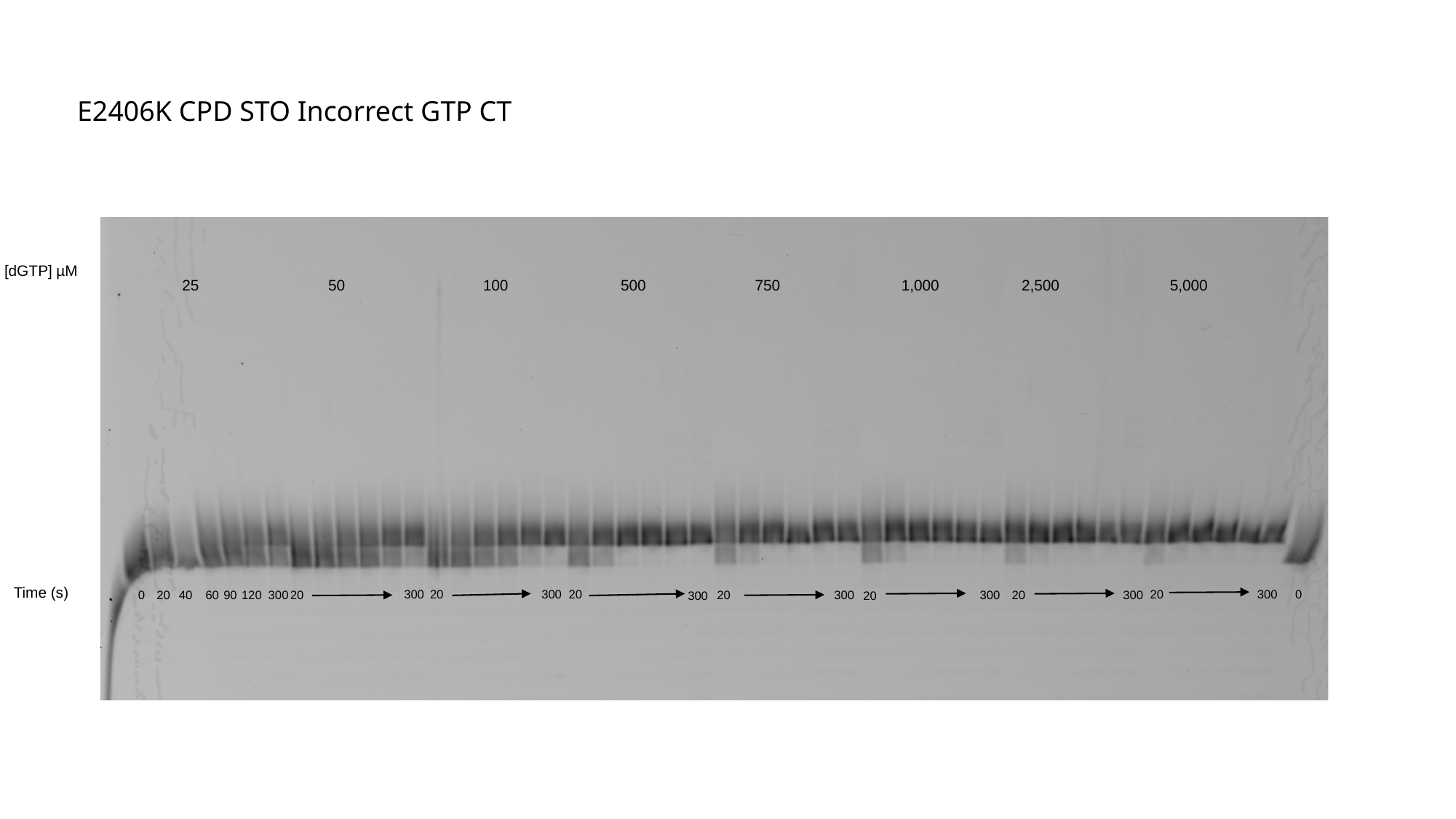

E2406K CPD STO Incorrect GTP CT
[dGTP] µM
50
100
500
750
1,000
2,500
5,000
25
Time (s)
300
20
300
20
300
0
20
0
20
40
60
90
120
300
300
20
300
20
300
20
300
20

### Slide 28
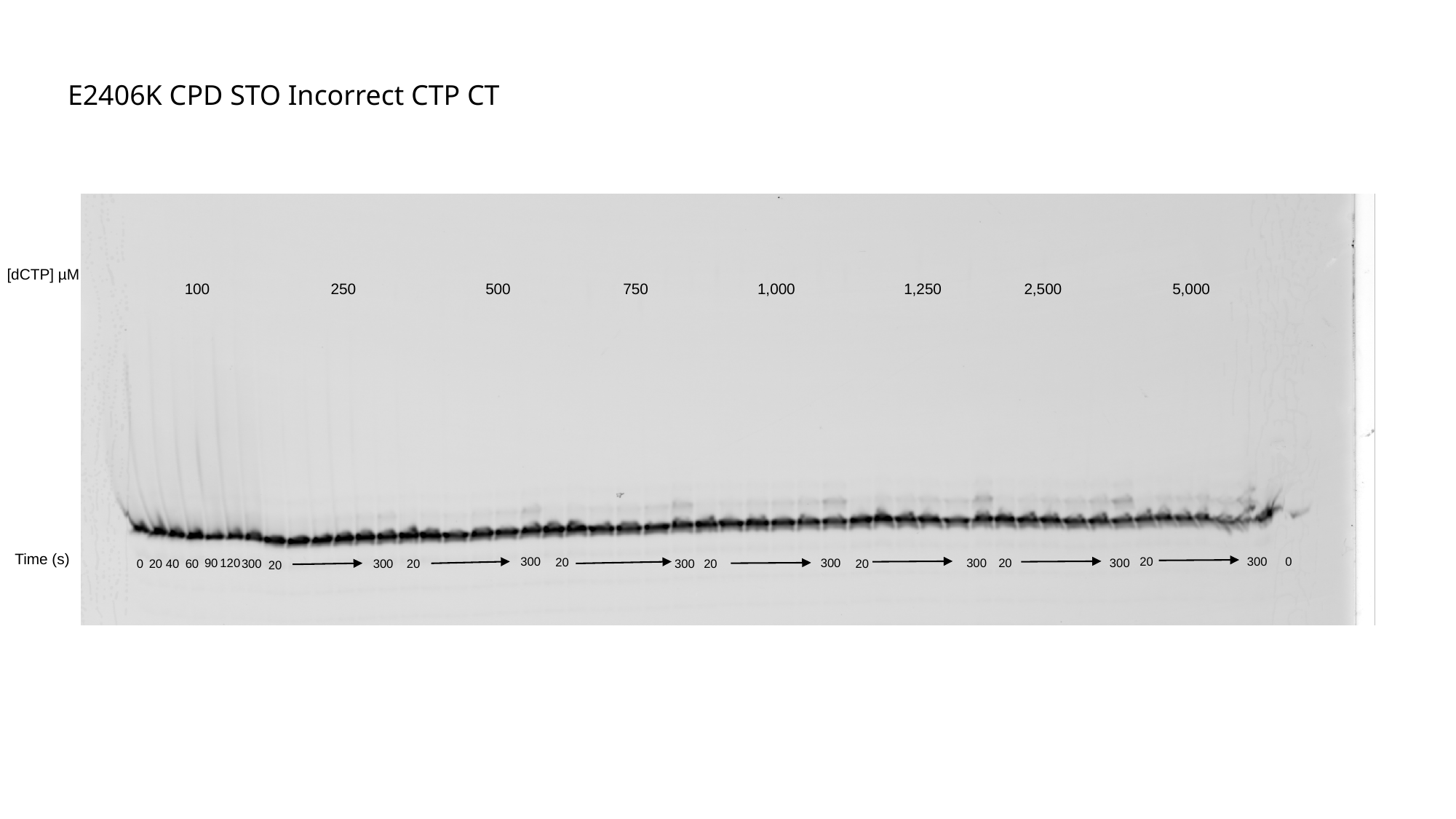

E2406K CPD STO Incorrect CTP CT
[dCTP] µM
250
500
750
1,000
1,250
2,500
5,000
100
Time (s)
300
20
300
0
20
300
20
300
300
90
120
20
0
20
40
60
300
300
20
300
20
20

### Slide 29
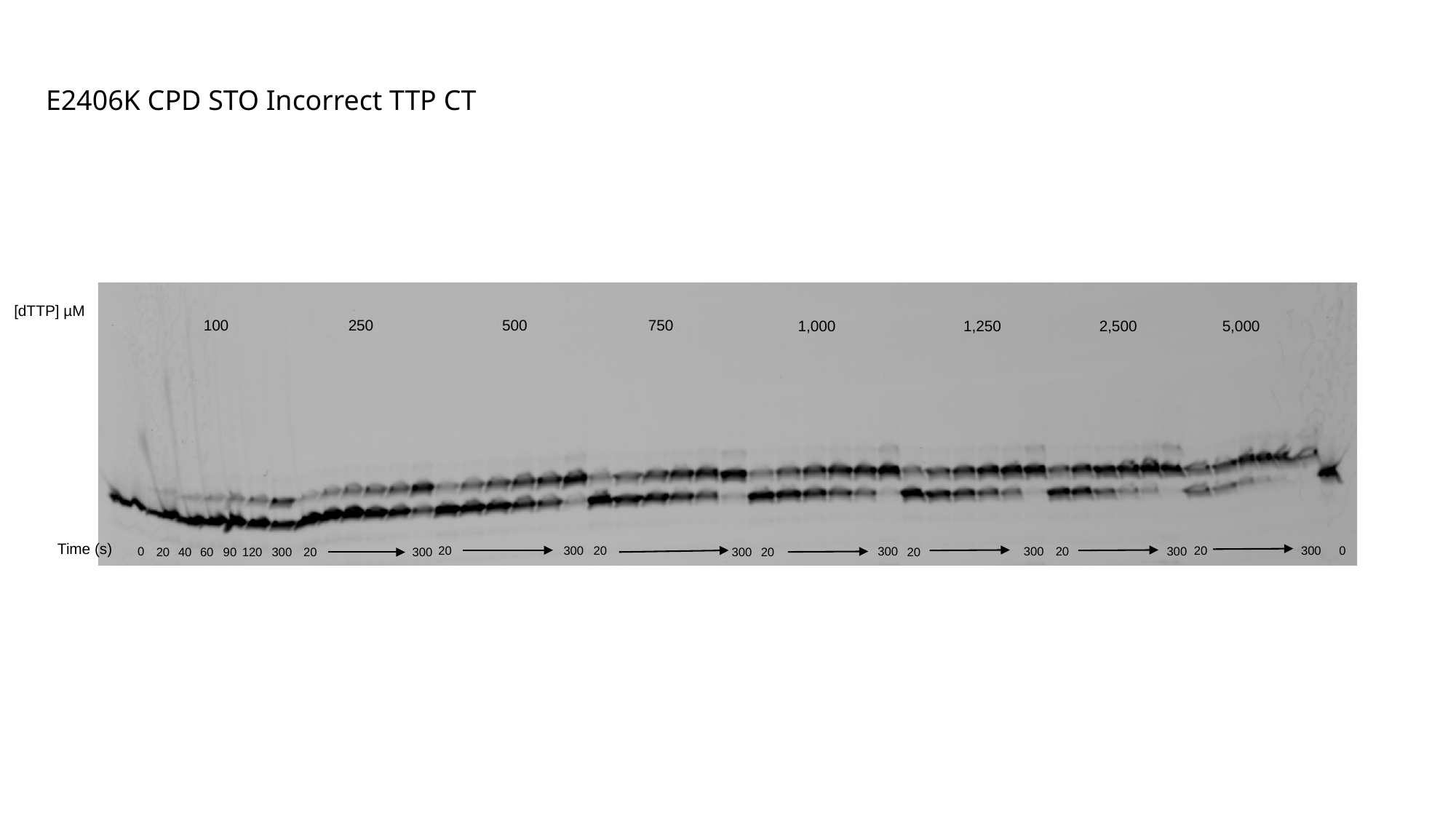

E2406K CPD STO Incorrect TTP CT
[dTTP] µM
100
250
500
750
1,000
1,250
2,500
5,000
Time (s)
20
300
20
20
300
0
0
300
20
300
300
20
300
20
40
60
90
120
300
20
300
20

### Slide 30
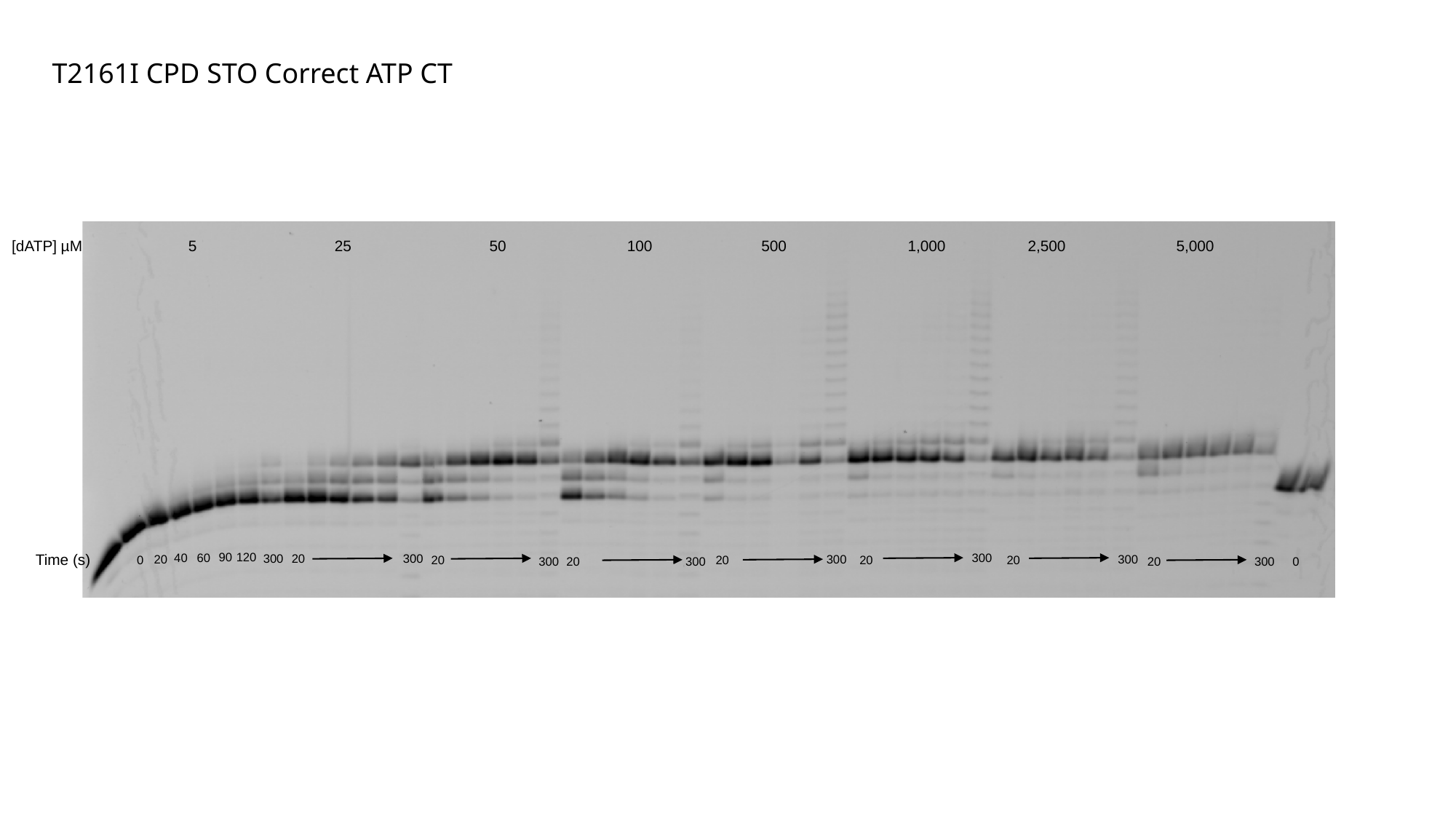

T2161I CPD STO Correct ATP CT
25
50
100
500
1,000
2,500
5,000
[dATP] µM
5
90
120
Time (s)
300
40
60
300
20
300
20
300
300
20
20
0
20
20
300
20
300
20
300
0

### Slide 31
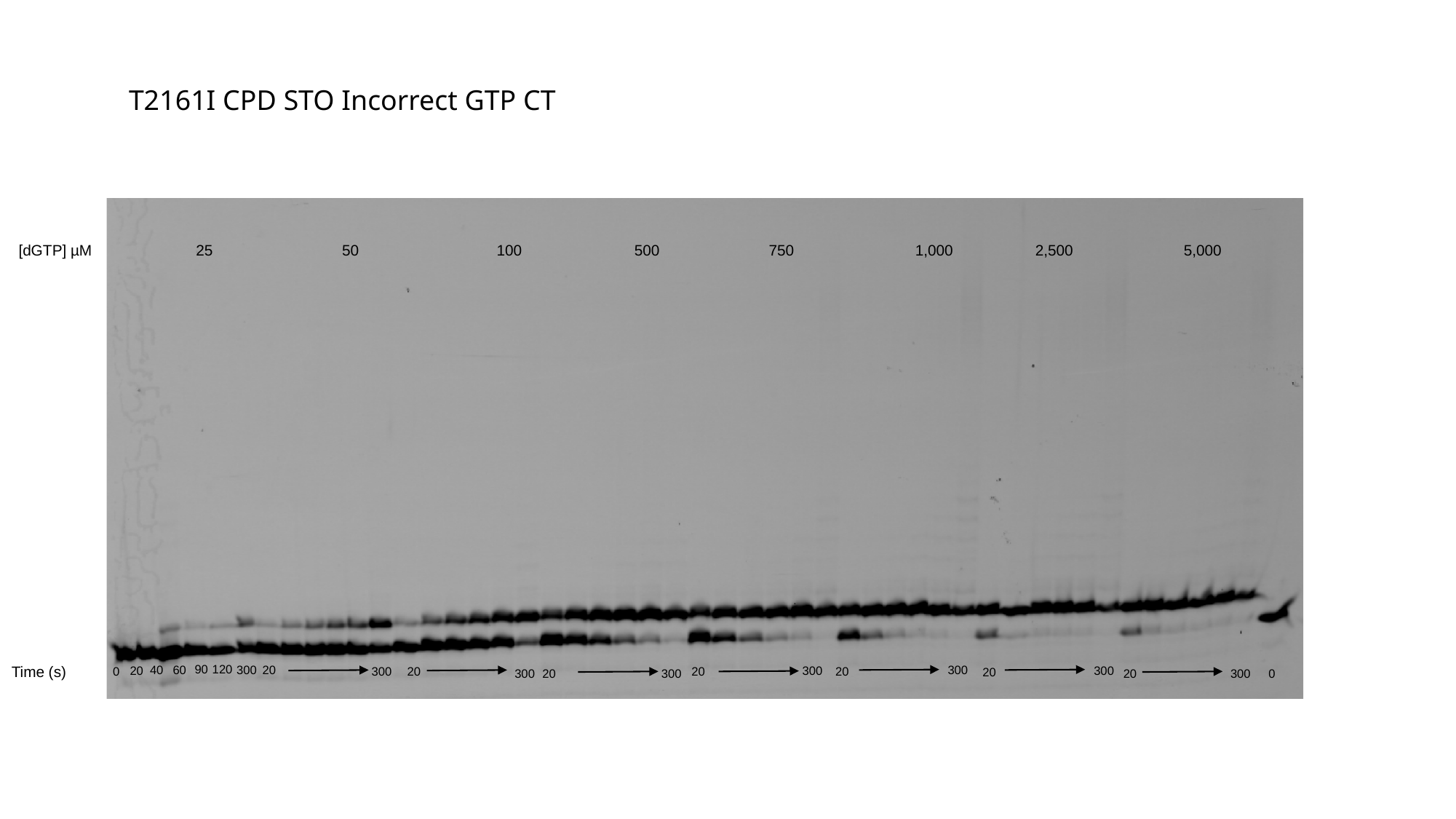

T2161I CPD STO Incorrect GTP CT
50
100
500
750
1,000
2,500
5,000
[dGTP] µM
25
90
120
Time (s)
300
40
60
300
20
20
300
300
300
20
20
0
20
20
300
20
300
20
300
0

### Slide 32
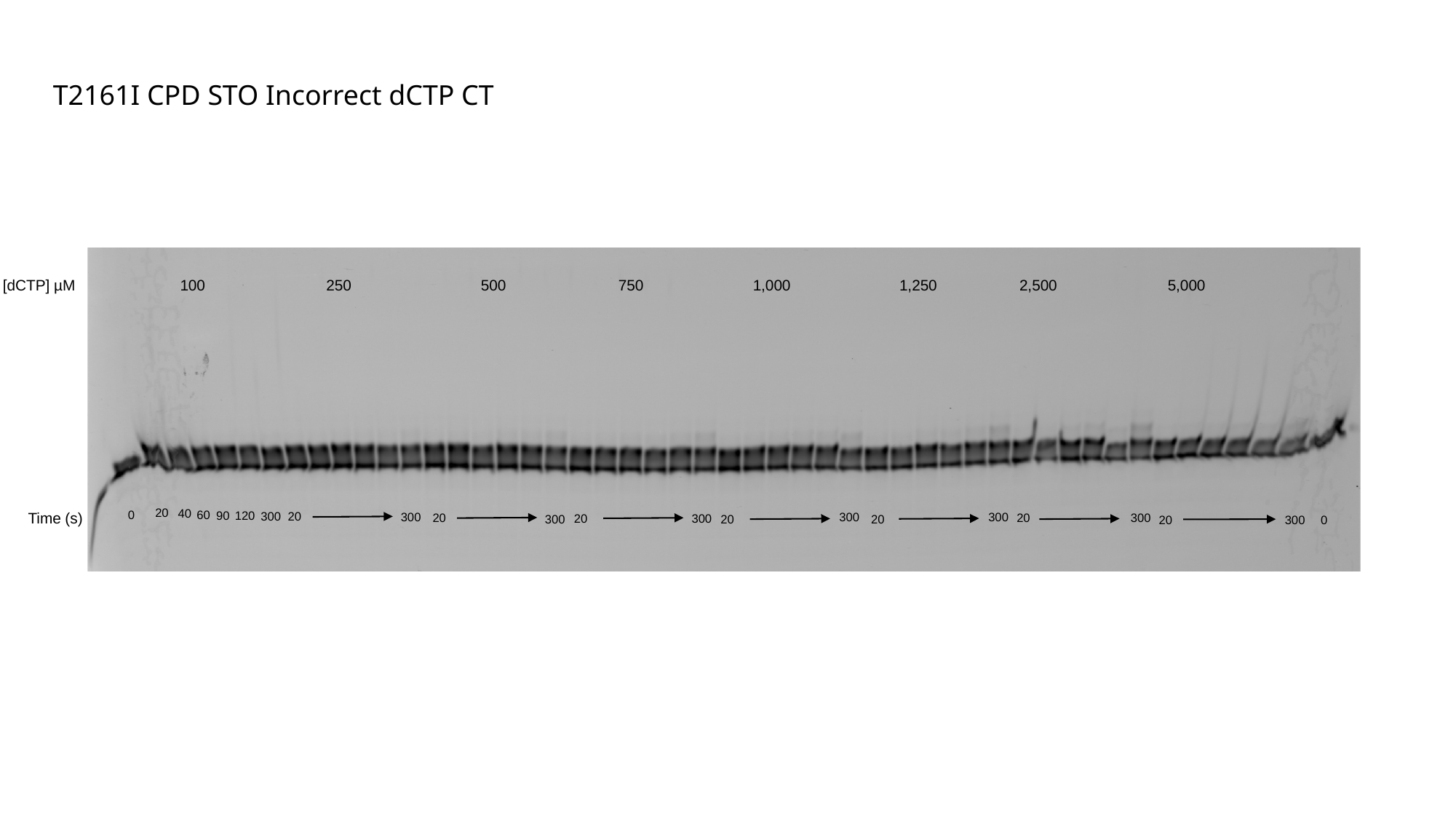

T2161I CPD STO Incorrect dCTP CT
250
500
750
1,000
1,250
2,500
5,000
[dCTP] µM
100
20
40
0
60
90
120
Time (s)
300
20
300
300
300
20
20
300
20
300
300
20
20
20
300
0

### Slide 33
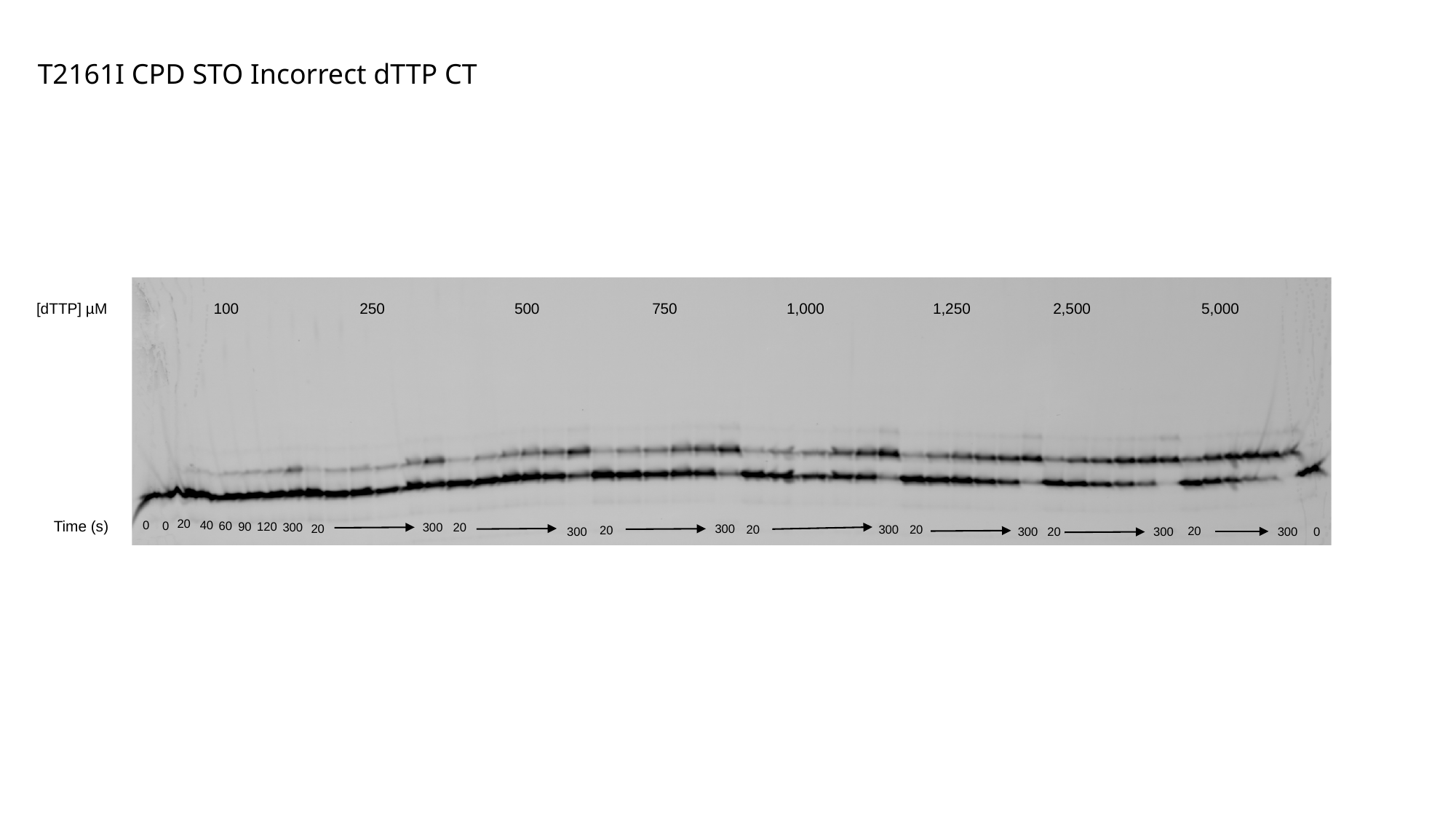

T2161I CPD STO Incorrect dTTP CT
250
500
750
1,000
1,250
2,500
5,000
[dTTP] µM
100
20
Time (s)
0
40
0
60
90
120
300
300
20
20
300
20
300
20
20
20
300
300
20
300
300
0
